# PilT and PilU are homohexameric ATPases that coordinate to retract type IVa pili

**DOI:** 10.1101/634048

**Authors:** Jennifer L. Chlebek, Hannah Q. Hughes, Aleksandra S. Ratkiewicz, Rasman Rayyan, Joseph Che-Yen Wang, Brittany E. Herrin, Triana N. Dalia, Nicolas Biais, Ankur B. Dalia

## Abstract

Bacterial type IV pili are critical for diverse biological processes including horizontal gene transfer, surface sensing, biofilm formation, adherence, motility, and virulence. These dynamic appendages extend and retract from the cell surface. In many type IVa pilus systems, extension occurs through the action of an extension ATPase, often called PilB, while optimal retraction requires the action of a retraction ATPase, PilT. Many type IVa systems also encode a homolog of PilT called PilU. However, the function of this protein has remained unclear because *pilU* mutants exhibit inconsistent phenotypes among type IV pilus systems and because it is relatively understudied compared to PilT. Here, we study the type IVa competence pilus of *Vibrio cholerae* as a model system to define the role of PilU. We show that the ATPase activity of PilU is critical for pilus retraction in PilT Walker A and/or Walker B mutants. PilU does not, however, contribute to pilus retraction in Δ*pilT* strains. Thus, these data suggest that PilU is a *bona fide* retraction ATPase that supports pilus retraction in a PilT-dependent manner. We also found that a Δ*pilU* mutant exhibited a reduction in the force of retraction suggesting that PilU is important for generating maximal retraction forces. Additional *in vitro* and *in vivo* data show that PilT and PilU act as independent homo-hexamers that may form a complex to facilitate pilus retraction. Finally, we demonstrate that the role of PilU as a PilT-dependent retraction ATPase is conserved in *Acinetobacter baylyi*, suggesting that the role of PilU described here may be broadly applicable to other type IVa pilus systems.

**Author Summary:** Almost all bacterial species use thin surface appendages called pili to interact with their environments. These structures are critical for the virulence of many pathogens and represent one major way that bacteria share DNA with one another, which contributes to the spread of antibiotic resistance. To carry out their function, pili dynamically extend and retract from the bacterial surface. Here, we show that retraction of pili in some systems is determined by the combined activity of two motor ATPase proteins.

## Introduction

Type IV pili are ubiquitous surface appendages in Gram-negative bacteria that promote diverse activities including attachment, virulence, biofilm formation, horizontal gene transfer, and twitching motility [1–5]. These structures can dynamically extend and retract from the cell surface, which is often critical for their function. A detailed mechanistic understanding of pilus dynamic activity, however, remains lacking.

Type IV pili are composed almost exclusively of a single protein called the major pilin, which forms a helical fiber that extends from the cell surface [6]. Extension and retraction of the pilus is hypothesized to occur through the interaction of cytoplasmic hexameric ATPases with the inner membrane platform of the pilus machine, whereby ATP hydrolysis likely facilitates turning of the platform in order to incorporate or remove major pilin subunits from the pilus fiber [7–9]. All type IV pilus systems encode a predicted ATPase, often called PilB, that facilitates pilus extension [7, 10–12]. Many type IVa pilus systems also contain a retraction ATPase, called PilT, which depolymerizes the pilus fiber and recycles pilin subunits into the inner membrane. Many type IVa pilus systems also encode another putative ATPase called PilU. PilU is a homolog of the retraction ATPase PilT, however, the function of this protein has remained unclear because PilU is relatively understudied compared to PilT and because of phenotypic inconsistency for *pilU* mutants in different organisms. For example, a *pilU* mutant of *Pseudomonas aeruginosa*, exhibits reduced twitching motility [13], but in other type IV pilus systems, including the *V. cholerae* competence pilus, loss of *pilU* does not reveal any overt phenotypes.

The facultative bacterial pathogen *Vibrio cholerae* uses a type IV competence pilus for DNA uptake during natural transformation. We have recently demonstrated that the type IV competence pili of *V. cholerae* can be fluorescently labeled by using a technique [14–16] in which an amino acid of the major pilin subunit PilA is replaced with a cysteine (PilA^S56C^ based on the mature pilin sequence, which was previously misannotated PilA^S67C^ [17] and PilA^S81C^ [18]), which allows for subsequent labeling with a fluorescently-labeled thiol-reactive maleimide dye (AF488-mal). This labeling approach does not impede dynamic pilus activity and allows for the direct observation and measurement of pilus extension and retraction by time-lapse epifluorescence microscopy. Using this labeling technique, we have recently demonstrated that the retraction of DNA-bound type IV competence pili is required to initiate the process of DNA uptake during natural transformation [18].

Here, we employ our pilus labeling approach and complementary molecular methods to address the role that PilU plays in type IV competence pilus retraction, and we extend our findings to the type IVa competence pilus of *Acinetobacter baylyi*. Our results indicate that PilT and PilU form independent homo-hexamers and that the ATPase activity of PilU promotes type IV pilus retraction in a PilT-dependent manner.

## Results

### PilU promotes type IV competence pilus retraction in a PilT-dependent manner

In order to study the function and role of ATPases in the retraction of type IV competence pili in *V. cholerae*, we utilize two main methods: (1) our labeling approach to directly measure pilus retraction via epifluorescence time-lapse microscopy and (2) assessing natural transformation, which is dependent on competence pilus retraction [18, 19]. Also, as for other pili [20–22], competence pili can mediate cell to cell interactions [17], which causes hyperpiliated strains to aggregate and pellet to the bottom of culture tubes. Consistent with prior work [19, 23–28], mutant strains that lack the retraction ATPase PilT are poorly transformable, exhibit markedly reduced rates of pilus retraction, aggregate, and are hyperpiliated (≥3 pili per cell) compared to the parent strain (Fig 1A, 1B, 1D and 1E). Also, the number of cells that exhibit dynamic pilus activity is reduced in the Δ*pilT* strain as observed by time-lapse epifluorescence microscopy, which is consistent with the hyperpiliated phenotype of this mutant (Fig 1C). Because the frequency of retraction was reduced by ∼2 logs in the Δ*pilT* strain (Fig 1C), we could not obtain enough events to make rigorous statistical comparisons. In the rare events observed, however, the retraction rate is reduced, albeit not significantly, ∼50-fold in the Δ*pilT* background.

**Fig 1.**
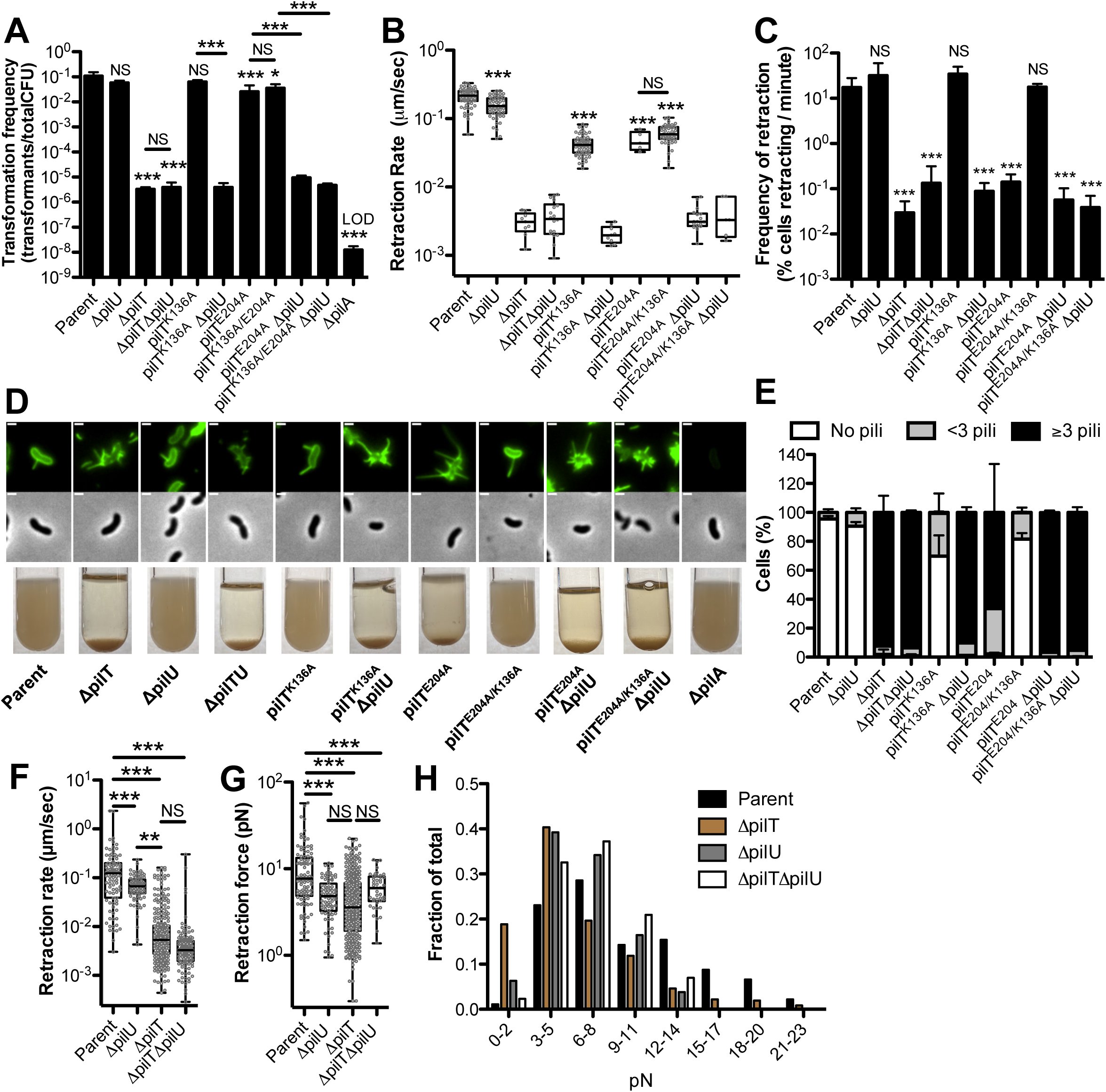
PilU mediates type IVa competence pilus retraction in a PilT-dependent manner in *V. cholerae*. **(A)** Natural transformation assays of the indicated strains. Reactions were incubated with 500 ng of transforming DNA overnight. Transformation frequency is reported as the fraction of cells that were transformed in each reaction (i.e. the number of transformants / total viable counts). Parent, *n* = 11, all other strains, *n* = 4. **(B)** Retraction rates of the indicated strains measured by epifluorescence time-lapse microscopy of AF488-mal labeled cells. Data are from three independent biological replicates; Parent, *n* = 68. Δ*pilU*, *n* = 56. Δ*pilT*, *n* = 9. Δ*pilT*Δ*pilU*, *n* = 18. *pilT^K136A^*, *n* = 66. *pilT^K136A^*Δ*pilU*, *n* = 9. *pilT^E204A^*, *n* = 5. *pilT^E204A/K136A^*, *n* = 50. *pilT^E204A^* Δ*pilU*, *n* = 17. *pilT^E204A/K136A^* Δ*pilU*, *n* = 6. **(C)** Frequency of retraction of the indicated strains measured by counting number of cells that exhibit a retraction event over a given time-lapse. Data are from three independent biological replicates; Parent, *n* = 1141. Δ*pilU*, *n* = 824. Δ*pilT*, *n* = 6681. Δ*pilT*Δ*pilU*, *n* = 3806. *pilT^K136A^*, *n* = 1048. *pilT^K136A^*Δ*pilU*, *n* = 997. *pilT^E204A^*, *n* = 4311. *pilT^E204A/K136A^*, *n* = 866. *pilT^E204A^* Δ*pilU*, *n* = 7243. *pilT^E204A/K136A^* Δ*pilU*, *n* = 6216. **(D)** Representative images of surface piliation (top) and aggregation phenotypes (bottom) for the indicated strains. Scale bar, 1 µm. **(E)** The percentage of cells with 0, <3 or ≥3 surface pili was determined from a static image. Data are from three independent biological replicates; Parent, *n* = 453. Δ*pilU*, *n* = 413. Δ*pilT*, *n* = 381. Δ*pilT*Δ*pilU*, *n* = 425. *pilT^K136A^*, *n* = 417. *pilT^K136A^* Δ*pilU*, *n* = 413. *pilT^E204A^*, *n* = 579. *pilT^E204A/K136A^*, *n* = 713. *pilT^E204A^* Δ*pilU*, *n* = 806. *pilT^E204A/K136A^* Δ*pilU*, *n* = 705. **(F)** Retraction rate (Parent, *n* = 95. Δ*pilU*, *n* = 83. Δ*pilT*, *n* = 324. Δ*pilT* Δ*pilU*, *n* = 125) and the **(G)** Box plot and **(H)** Histogram of the retraction force (Parent, *n* = 101. Δ*pilU*, *n* = 90. Δ*pilT*, *n* = 372. Δ*pilT* Δ*pilU*, *n* = 43) was measured via micropillar assays. All bar graphs are shown as the mean ± SD. All Box plots represent the median and the upper and lower quartile, while the whiskers demarcate the range. Asterisk(s) directly above bars denote comparisons to the parent strain. All comparisons were made by one-way ANOVA with Tukey’s post test. LOD, limit of detection; NS, not significant; ** = *P* < 0.01, *** = *P* < 0.001.

The residual retraction in a Δ*pilT* strain is not due to PilU because a Δ*pilT* Δ*pilU* double mutant strain displays the same phenotype as a Δ*pilT* strain (Fig 1A-E). By contrast, Δ*pilU* mutants are highly transformable, do not aggregate, exhibit normal numbers of pili and are largely indistinguishable from the parent strain (Fig 1A and 1C-E) aside from a significant 1.3-fold reduction in retraction rates compared to the parent strain (Fig 1B). This effect of PilU on retraction rate is much smaller compared to the ∼50-fold reduction in retraction rate observed in the Δ*pilT* and Δ*pilT* Δ*pilU* mutants (Fig 1B). This suggests that PilT can still mediate retraction in a Δ*pilU* strain (because retraction frequency and rates for Δ*pilU* are closer to the parent than the Δ*pilT* Δ*pilU* strain), while PilU cannot mediate retraction in a Δ*pilT* strain (since the Δ*pilT* strain resembles the Δ*pilT* Δ*pilU* strain). Thus, these data support a model where PilT plays the dominant role in pilus retraction, which is consistent with prior reports [18, 19, 26].

PilT is a predicted ATPase and its ATPase activity has been shown to be important in the function of other type IV pilus systems [12, 29]. Therefore, we hypothesized that the ATPase activity of PilT would be required for retraction of the competence pilus. In order to answer this question, we putatively ablated the ATPase activity of PilT by introducing a mutation in either an invariantly conserved lysine in the Walker A motif required for ATP binding (*pilT^K136A^*), a mutation in a conserved glutamic acid in the Walker B motif required for ATP hydrolysis (*pilT^E204A^*), or by making a double mutant containing both mutations (*pilT^E204A/K136A^*). Though these Walker A and Walker B mutations have not been well studied in *V. cholerae* PilT-like ATPases, the effect of these mutations has been well characterized in other systems and we assume that they ablate the ATPase activity of this protein in a similar manner [30]. Interestingly, we found that strains harboring mutations in PilT that putatively ablate the ATPase activity of this protein (*pilT^K136A^*, *pilT^E204A^*, *pilT^E204A/K136A^*) all transform similarly to the parent strain and retract competence pili at rates that are at most ∼4-fold lower than the parent (Fig 1A-B). This is in stark contrast to a Δ*pilT* strain where the transformation frequency is reduced ∼4-logs and retraction is >50-fold slower compared to the parent strain (Fig 1A-B). These results were quite surprising and suggested that either (1) the ATPase activity of the retraction machinery is not critical for promoting pilus retraction, (2) that the mutations to the Walker motifs do not ablate PilT ATPase activity, or (3) that another retraction ATPase may promote retraction in the absence of PilT ATPase activity.

We tested the third model and specifically assessed whether PilU could promote retraction in predicted PilT ATPase-deficient mutants. Indeed, we found that *pilT^K136A^*Δ*pilU*, *pilT^E204A^*Δ*pilU* and *pilT^E204A/K136A^*Δ*pilU* strains all display reduced transformation, are hyperpiliated, aggregate, exhibit markedly reduced retraction frequency, and reduced retraction rates similar to a Δ*pilT* Δ*pilU* strain (Fig 1A-E). This is consistent with PilU promoting retraction in the *pilT^K136A^, pilT^E204A^* and *pilT^E204A/K136A^*backgrounds. Importantly, the *pilT^K136A^*Δ*pilU*, *pilT^E204A^*Δ*pilU* and *pilT^E204A/K136A^*Δ*pilU* strains could be complemented via ectopic expression of *pilT* or *pilU* (S1 Fig). Since loss of *pilT* alone (i.e. Δ*pilT*) results in a phenotype comparable to a Δ*pilT* Δ*pilU* mutant, this result indicates that PilU is not sufficient to drive pilus retraction in the absence of PilT (Fig 1A-E). Consistent with this, ectopic expression of *pilU* does not rescue the transformation deficit of a Δ*pilT* Δ*pilU* mutant (S1 Fig). Together, these data suggest that PilU supports pilus retraction in a PilT-dependent manner.

Interestingly, whereas a *pilT^K136A^*Walker A mutant displays normal piliation, the *pilT^E204A^*Walker B mutant is hyperpiliated (Fig 1D-E). The frequency of cells that exhibit pilus retraction in the *pilT^E204A^*mutant is much lower than the parent and comparable to the Δ*pilT* Δ*pilU* strain (Fig 1C), which may explain this hyperpiliated phenotype; however, when pilus retraction is observed, the rate of retraction is closer to that of the parent than the Δ*pilT* Δ*pilU* mutant (Fig 1B). Considering the canonical functions of the Walker A and B motifs in ATP-binding and hydrolysis, respectively, the *pilT^E204A^* Walker B mutant should be able to bind to ATP but not hydrolyze it. Because the putative ATP-binding deficient *pilT^K136A^*Walker A mutant exhibited parental levels of piliation (Fig 1D-E), we hypothesized that the hyperpiliation observed in *pilT^E204A^*may be due to the fact that this protein may be locked in an ATP-bound state. A recent preprint shows that different ATP-bound states of PilT exhibit different conformations [31]. This may suggest that the conformation adopted by PilT^E204A^ reduces the ability of PilU to facilitate retraction. Because ATP-binding must precede hydrolysis, we hypothesized that a Walker A and B double mutant (*pilT^E204A/K136A^*) should restore the ability of PilU to mediate normal piliation because this would prevent the formation of the ATP-bound conformation of PilT. Consistent with this, we observed that the phenotype of the putative ATP-binding deficient Walker A mutation (*pilT^K136A^*) is dominant over the putative ATP-hydrolysis deficient *pilT^E204A^* Walker B mutation (Fig 1D-E). Both the Walker A and Walker B mutations should result in loss of ATPase activity. Therefore, to avoid phenotypic consequences associated with the ATP-bound state of the *pilT^E204A^* mutant, we utilized the *pilT^K136A^* mutant as a putative ATPase-defective allele of PilT moving forward.

### PilU is required for forceful retraction

The data thus far suggest that PilU is capable of supporting pilus retraction in putative PilT ATPase-deficient mutants, but that PilU is not necessary to mediate pilus retraction (since Δ*pilU* mutants transform at frequencies that are similar to the parent). PilU, however, is present in many type IVa pilus systems, so the question remains: Why is PilU so widely conserved if it is not necessary for pilus retraction? Recent work in *Pseudomonas aeruginosa* indicates that PilU may be required for forceful retraction [32]. Our data thus far indicate that the Δ*pilU* mutant only causes a modest reduction in the retraction rate (Fig 1A), which does not have a phenotypic consequence on pilus-dependent DNA uptake (Fig 1C); however, these measurements are based on free pili that are not retracting under load. We hypothesized that PilU might be required for forceful retraction, such as when pili are attached to a surface. To test this, we assessed pilus retraction using a micropillar assay in which binding of pili to micropillars and subsequent retraction results in measurable bending and displacement of the micropillars.

These data can then be used to calculate the rates and force of pilus retraction [33]. In agreement with our cell biological approach (i.e. pilus labeling), the micropillar assay showed that retraction rates were reduced in both Δ*pilU* and Δ*pilT* strains; with PilT having a larger effect on this rate (2-fold reduction for Δ*pilU* vs 14-fold reduction for Δ*pilT*) (Fig 1B and 1F).

Importantly, the micropillar assay is more sensitive at detecting rare pilus retraction, which allowed us to collect enough events to show that Δ*pilT* mutants are statistically significantly reduced for retraction rates compared to the parent strain, which is consistent with previous work [18]. In contrast, the Δ*pilU* and Δ*pilT* strains exhibited a similar reduction in the retraction force (∼2-fold for both) compared to the parent strain (Fig 1G). As other groups have also reported, data collected on pilus retraction forces tend to be highly variable [18, 32, 34], which we also observed here (Fig 1G-H). Despite this variability, a histogram reveals that PilT and PilU promote retraction forces >12 pN because retraction events exceeding this force are relatively rare in the Δ*pilT* and Δ*pilU* strains (Fig 1H). Thus far, we have shown that Δ*pilU* and Δ*pilT* mutants display distinct phenotypes for transformation frequency (Fig 1A), retraction rates (Fig 1B and 1F) and piliation state (Fig 1D-E). Interestingly, the micropillar data reveal that Δ*pilU* and Δ*pilT* mutants have a similar effect on retraction force (Fig 1G-H). This suggests that PilU and PilT are both necessary for generating maximal retraction forces in the *V. cholerae* competence pilus.

### The ATPase activities of PilT and PilU define the rate of type IV competence pilus retraction

Just like PilT, PilU is a PilT-like ATPase with a conserved lysine in its Walker A motif. Because we observed that PilU supports pilus retraction in the *pilT^K136A^* mutant (Fig 1B), we hypothesized that the ATPase activity of PilU may be important for supporting pilus retraction. To test the relative roles of each of these ATPases in pilus retraction, we mutated the conserved lysine in the Walker A motifs of PilT and/or PilU. The *pilU^K134A^* strain retracts ∼1.5-fold slower than the parent strain, similar to the Δ*pilU* mutant (Fig 2A). There may be a limited effect of the *pilU^K134A^* mutation on pilus retraction because this strain still has a functional PilT ATPase. However, we hypothesized that PilU ATPase activity may be critical for retraction in a background where PilT ATPase activity is also inactivated. Consistent with this, we found that a *pilT^K136A^ pilU^K134A^* strain is indistinguishable from a *pilT^K136A^*Δ*pilU* strain for pilus retraction, piliation, aggregation, and transformation (Fig 2A-C). Together, these data suggest that PilU ATPase activity promotes type IV competence pilus retraction.

**Fig 2.**
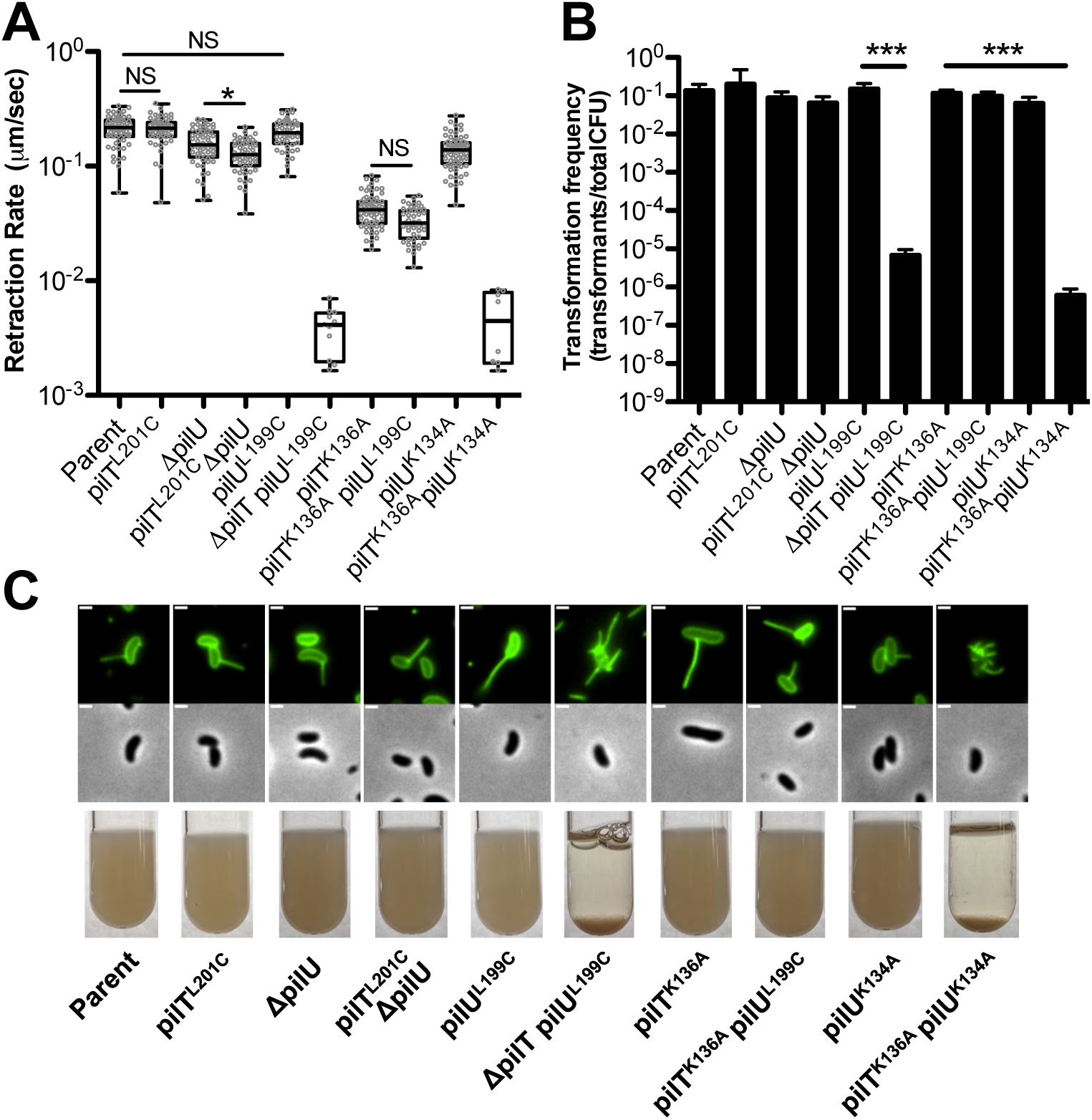
The ATPase activity of PilT and PilU mediate pilus retraction. **(A)** Retraction rates of the indicated strains measured by epifluorescence time-lapse microscopy of AF488-mal labeled cells. Box plots represent the median and the upper and lower quartile, while the whiskers demarcate the range. Data are from three independent biological replicates; *pilU^K134A^*, *n* = 70. *pilT^K136A^ pilU^K134A^*, *n* = 8. *pilT^K136A^ pilU^L199C^*, *n* = 46. *pilU^L199C^*, *n* = 49. *pilT^L201C^*, *n* = 54. *pilT^L201C^* Δ*pilU*, *n* = 54. Δ*pilT pilU^L199C^*, *n* = 11. Data for the retraction rates of the parent, Δ*pilU,* and *pilT^K136A^* strains are identical to those shown in Fig 1B and plotted again here to facilitate comparisons to the mutant strains indicated. **(B)** Natural transformation assays of the indicated strains incubated with 500 ng of transforming DNA overnight. Data are shown as the mean ± SD. Parent, *n* = 6. All other strains, *n* = 4**. (C)** Representative images of surface piliation (top) and aggregation phenotypes (bottom) for the indicated strains. Scale bar, 1 µm. Asterisk(s) directly above bars denote comparisons to the parent strain. All comparisons were made by one-way ANOVA with Tukey’s post test. NS, not significant; * = *P* < 0.05, *** = *P* < 0.001.

The data above suggest that the ATPase activities of PilT and PilU are critical for pilus retraction. These ATPases are predicted to be motor proteins that facilitate the depolymerization of pilus fibers. Thus, if we alter the ATPase rates of these motors, we hypothesized that this should alter the rates of pilus retraction. To test this hypothesis, we mutated a leucine in the Walker B domain of PilT and PilU to a cysteine, which was previously shown to reduce the rate of ATP hydrolysis of *Neisseria gonorrhoeae* PilT ∼2-fold [26]. When these Walker B mutations were tested in isolation, they did not alter the rate of pilus retraction (comparing the parent strain to *pilT^L201C^* and *pilU^L199C^*) (Fig 2A). This may be due to the fact that these cells still have one fully functional retraction ATPase (i.e. *pilT^L201C^* still has a fully functional PilU and *pilU^L199C^* still has a fully functional PilT). We hypothesized that these putatively Walker B “slow” ATPase mutations, however, may have an impact on pilus retraction if they represented the sole functional retraction ATPase in the cell. To that end, we generated both a *pilT^L201C^*Δ*pilU* strain and a *pilT^K136A^ pilU^L199C^* strain. The *pilT^L201C^* Δ*pilU* mutant showed a slight (1.3 fold), but significant, reduction in retraction rate compared to the Δ*pilU* strain (Fig 2A), which provides genetic evidence to suggest that the ATPase rate of PilT may be linked to the retraction rate of the type IV competence pilus and further supports the idea that PilT promotes pilus retraction through its ATPase activity. The *pilT^K136A^ pilU^L199C^*mutant also showed a slight (1.5-fold) reduction in retraction rate compared to *pilT^K136A^*, although this difference was not statistically significant (Fig 2A). Because this difference was not statistically significant, it may suggest that the ATPase rate of PilU does not dictate the retraction rate of competence pili even when it is the sole retraction ATPase; however, because this mutant trended towards a decreased retraction rate, this result may still suggest that the ATPase rate of PilU is linked to retraction rates.

### PilT and PilU directly interact with one another

All of these data presented above indicate that PilT and PilU can both support pilus retraction. Therefore, we hypothesized that PilT and PilU may interact to form a complex that supports pilus retraction. To address this, we utilized a Bacterial Adenylate Cyclase Two-Hybrid (BACTH) assay [35, 36]. As expected for predicted hexameric proteins like PilT and PilU, the BACTH analysis showed PilT-PilT and PilU-PilU self-interactions (Fig 3 and S2 Fig). Additionally, the BACTH assay showed that PilT and PilU interact with one another, which is consistent with prior work in other systems [35, 37], and further demonstrated that Walker A mutations in either protein (PilT^K136A^ and PilU^K134A^) did not disrupt this interaction (Fig 3 and S2 Fig). Thus, our BACTH results suggest that PilT and PilU may form a complex. These results do not, however, indicate whether PilU and PilT form heteromeric complexes or homomeric complexes to facilitate pilus retraction. We test this further below.

**Fig 3.**
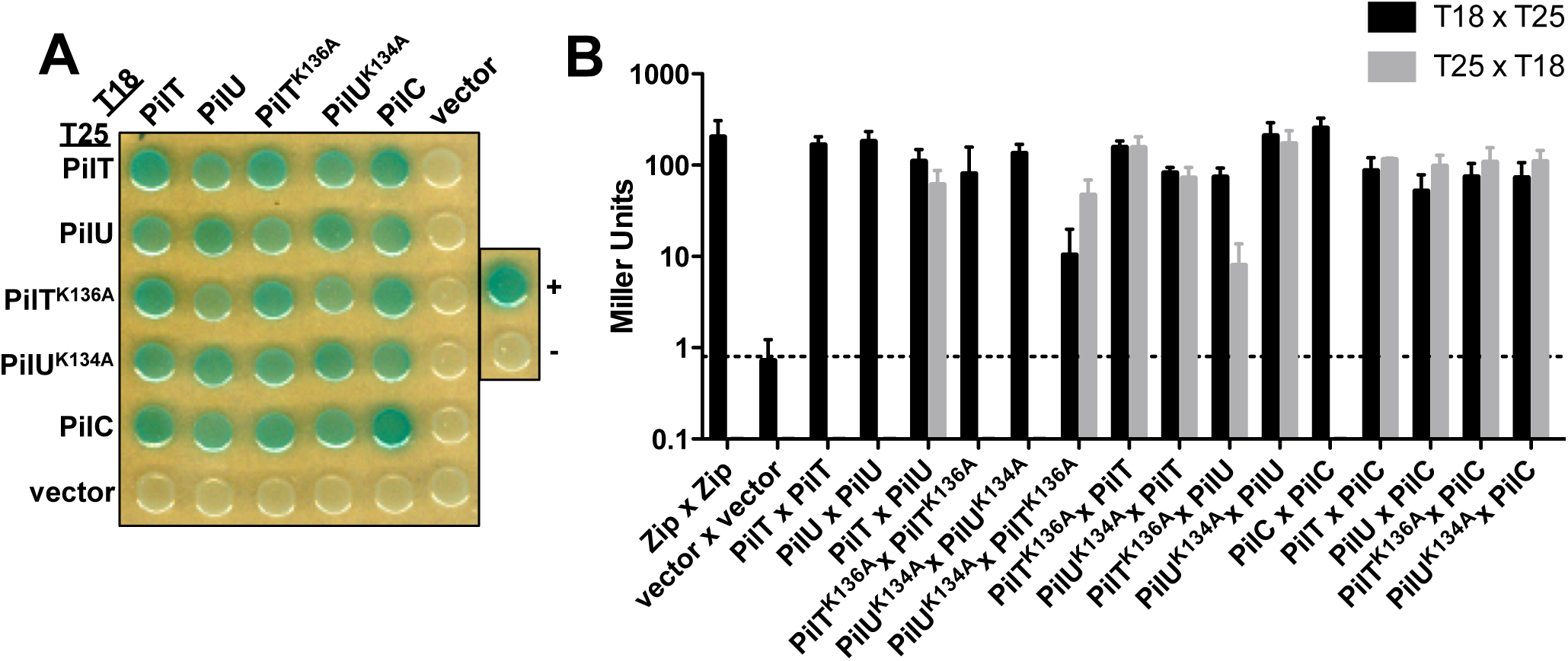
PilT and PilU interact with each other and the pilus platform protein PilC. Bacterial Adenylate Cyclase Two-Hybrid (BACTH) assays were performed to test pair-wise interactions between full length PilT, PilU, PilT^K136A^, PilU^K134A^, and PilC that were N-terminally fused to the T18- and T25-fragments of adenylate cyclase. Each construct was also tested for interactions with empty T18 and T25 vectors as a negative control. Leucine zipper fusions to T18 and T25 were used as the positive control (+) and empty T18 and T25 vectors served as an additional negative control (-). **(A)** Representative image of the BACTH assay plated on X-gal-containing media. **(B)** BACTH miller assays of the indicated pairwise interactions. The dotted line represents the β-galactosidase activity of the empty vector negative control. Data are shown as the mean ± SD and are from three independent experiments. Statistical analysis of these data can be found in S2 Fig.

### PilT and PilU form independent homohexameric rings *in vitro* and *in vivo*

PilT in other systems has been shown to form a hexameric complex [38], and by homology we hypothesized that PilU also forms a hexamer. Based on transformation frequency assays, N-terminally tagged ^6X^His-PilT and ^6X^His-PilU were fully functional (S3A Fig). Therefore, we used our purified ^6X^His-PilT and ^6X^His-PilU to characterize these proteins *in vitro*. Negative stain transmission electron microscopy revealed that both ^6X^His-PilT and ^6X^His-PilU formed hexameric ring complexes (Fig 4A). These results indicate that these proteins are capable of forming independent homohexameric complexes *in vitro*. PilT and PilU have both sequence (∼65% similarity) and structural similarity to one another (Fig 4A), thus it is also possible that these proteins can be incorporated into one ring and form heterohexameric complexes that facilitate pilus retraction *in vivo*.

**Fig 4.**
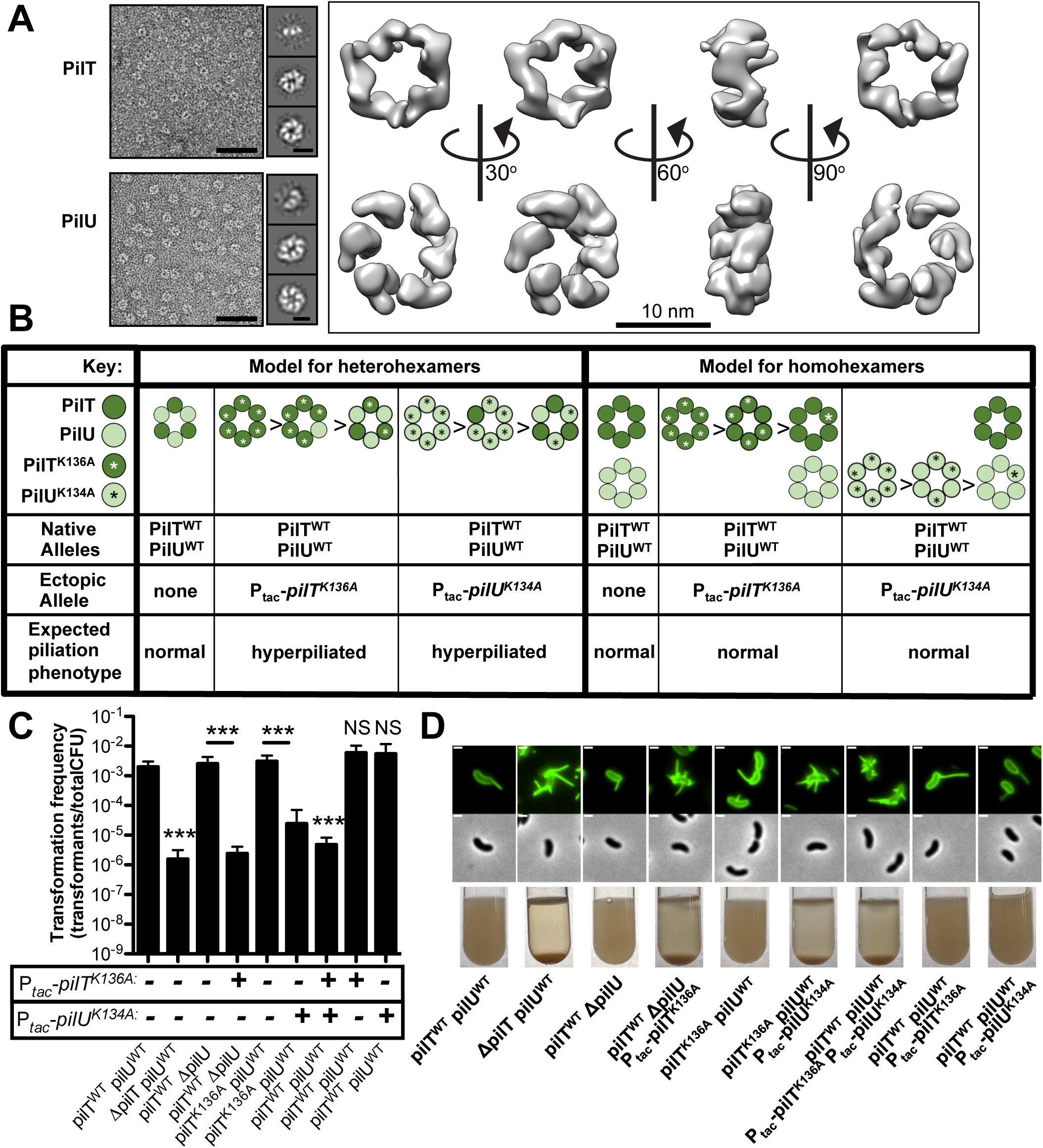
PilT and PilU form independent homomeric complexes *in vitro* and *in vivo* to facilitate pilus retraction. **(A)** Representative negative stain transmission electron micrographs of purified ^6X^His-PilT and ^6X^His-PilU (left; scale bar, 50 nm). 2D class averages revealing hexameric rings (middle; scale bar, 10 nm) and 3D reconstructions (right) of the ring complex. **(B)** Schematic of the experimental approach and expected outcomes for testing whether PilT and PilU form homomeric or heteromeric complexes to facilitate pilus retraction *in vivo*. **(C)** Natural transformation assays of indicated strains incubated with 500 ng of tDNA and after 7 minutes, DNAseI was added to prevent additional DNA uptake. The genotype of native *pilT* and *pilU* alleles are written below each bar graph and the presence or absence of a chromosomally-integrated IPTG-inducible ectopic allele of *pilT^K136A^* or *pilU^K134A^* is indicated by a + or -. Strains were grown with 100 µM IPTG if they contained an ectopic expression construct. Data are shown as the mean ± SD and are from four independent biological replicates. **(D)** Representative images of surface piliation (top) and aggregation phenotypes (bottom) for the indicated strains. Scale bar, 1 µm. Asterisk(s) directly above bars denote comparisons to the parent strain. All comparisons were made by one-way ANOVA with Tukey’s post test. NS, not significant; *** = *P* < 0.001.

In order to test if PilT and PilU form heterohexamers or independent homohexamers to facilitate pilus retraction *in vivo*, we hypothesized that overexpression of putative ATPase-inactive mutants (i.e. PilT^K136A^ and PilU^K134A^) in backgrounds that have native PilT and PilU would allow us to distinguish between these two models. For example, if these proteins form heteromeric complexes, then we would predict that overexpression of either putative ATPase-inactive protein (PilT^K136A^ or PilU^K134A^) should result in complexes that are mostly composed of ATPase-inactive proteins and result in inhibition of pilus retraction (yielding a hyperpiliated phenotype) and reduce rates of natural transformation (Fig 4B). If these proteins solely form homomeric complexes, however, then we would predict that overexpression of only one ATPase-inactive protein (PilT^K136A^ or PilU^K134A^) should only inhibit the activity of one motor complex (PilT or PilU, respectively) and should not have a dramatic effect on pilus retraction or natural transformation because the other retraction motor would still be intact (Fig 4B).

In order for us to successfully test this question, we needed to: (1) be able to ectopically overexpress ATPase deficient alleles of PilT or PilU, (2) demonstrate that ATPase deficient alleles can interact with their ATPase sufficient counterparts and form proper hexamers, and (3) show that ectopic expression of ATPase deficient alleles of PilT or PilU can inhibit the activity of their respective natively expressed ATPase. We test each of these in turn below. First, we confirmed that we could ectopically overexpress PilT and PilU. Using functional 3xFLAG-tagged versions of PilT and PilU (S3A-B Fig), we show that our chromosomal *P_tac_*regulated constructs resulted in dramatic overexpression of these proteins (S3C Fig). Importantly, overexpression of WT PilT and PilU did not affect pilus activity or natural transformation (S1, S3B **and** S3D Figs). Next, we purified 6xHis-PilT^K136A^ and 6xHis-PilU^K134A^ and demonstrated that they formed ring structures *in vitro* (S3E Fig), indicating that these proteins should form complexes with ATPase sufficient alleles. This is also supported by our BACTH data (Fig 3). Finally, we wanted to confirm that overexpressing the ATPase-inactive PilT^K136A^ or PilU^K134A^ could effectively impede the function of the natively expressed ATPase-active PilT or PilU. To test this, we genetically isolated either PilT or PilU as the sole functional retraction ATPase (i.e. PilT activity is isolated in a Δ*pilU* background, while PilU activity is isolated in a *pilT^K136A^* background) and then overexpressed the cognate ATPase-inactive allele (P_tac_-*pilT^K136A^* or P_tac_-*pilU^K134A^*, respectively). We assessed piliation, aggregation, and natural transformation as indicators of pilus retraction. We found that, as expected, overexpression of ATPase-inactive alleles was sufficient to disrupt the activity of the natively expressed ATPase (Fig 4C-D -compare Δ*pilU* to Δ*pilU P_tac_*-*pilT^K136A^*and *pilT^K136A^* to *pilT^K136A^ P_tac_*-*pilU^K134A^*). Also, as expected, overexpression of both ATPase-deficient alleles together (P_tac_-*pilT^K136A^*and P_tac_-*pilU^K134A^*) in a background that has natively expressed PilT and PilU, results in hyperpiliation, aggregation and reduced natural transformation (Fig 4C-D), which further demonstrates that overexpression of these ATPase-deficient proteins inhibits the activity of their natively expressed ATPase–sufficient versions.

The experiments above indicated that it would be feasible to test whether PilT and PilU formed homomeric or heteromeric complexes *in vivo* as described in our model (Fig 4B). When we overexpressed only one ATPase-deficient protein (P_tac_-*pilT^K136A^*or P_tac_-*pilU^K134A^*) in a background that has natively expressed PilT and PilU, we found that piliation and transformation were indistinguishable from the parent (Fig 4C-D). Thus, these data are consistent with PilT and PilU forming independent homohexamers to facilitate pilus retraction.

Using our purified PilT and PilU, we also attempted to directly measure the ATPase activity of these proteins. Unfortunately, the ATPase activity of these purified proteins was indistinguishable from contaminating ATPases and/or below our detection limit (∼1 nmol pi/min/mg protein). Poor ATPase activity in purified retraction ATPases is not unique to our system [12, 29, 39]. Alternatively, these results may suggest that the ATPase activities of these proteins are stimulated *in vivo* via interaction with the proteins that compose the pilus machine, as previously suggested [12]. Regardless, our previous *in vivo* data (Fig 1A-D) are consistent with the *pilT^K136A^*mutant being ATPase deficient.

### The presence of PilT is not required for PilU to interact with the pilus platform PilC

Our data thus far indicate that PilU requires the presence of PilT (whether it is a functional ATPase or not) to support optimal pilus retraction. The extension and retraction ATPases are hypothesized to interact with the platform protein PilC to mechanically facilitate polymerization and depolymerization of the pilus fiber [7, 37, 40]. Because PilU is dependent on the presence of PilT to mediate retraction, we hypothesized that PilT can interact with PilC, but that PilU cannot. To test this, we utilized the BACTH assay to test the interactions of PilT and PilU with PilC. This assay showed that PilT and PilU could both interact with PilC, and that the Walker A mutations of these ATPases (PilT^K136A^ and PilU^K134A^) did not affect these interactions (Fig 3). It is necessary to point out, however, that the BACTH assay cannot address relative affinities of protein interactions due to non-native expression levels, but instead indicates the potential presence or absence of an interaction. The biological relevance of these interactions will depend on the relative affinities and concentrations of these proteins in their natural context. The interaction between PilC and PilT supports the canonical model that retraction ATPases interact with PilC to facilitate depolymerization of the pilus fiber. Interestingly, our data suggest that PilC and PilU interact independently of PilT. This result is also supported by prior BACTH analysis of *P. aeruginosa* pilus components [37]. Thus, the simple model of PilU requiring PilT for interaction with the platform PilC appears to be incorrect.

### The function of PilU as a PilT-dependent retraction ATPase is conserved in the Acinetobacter baylyi competence pilus

As mentioned earlier, many type IVa pilus systems encode PilU in addition to the canonical retraction ATPase PilT. Therefore, we sought to assess whether PilU could facilitate PilT-dependent retraction in other type IV pilus systems. To test this, we studied the type IVa competence pili of a distinct gammaproteobacterium *Acinetobacter baylyi*. *A. baylyi* competence pili also mediate DNA-uptake and the genetic organization of *pilT* and *pilU* is similar to that observed in *V. cholerae*.

The competence pili of *A. baylyi* likely facilitate DNA uptake in a retraction-dependent manner, similar to the competence pili of *V. cholerae*. Thus, we employed natural transformation assays to dissect the role that PilT and PilU play in *A. baylyi* competence pilus retraction. These assays showed that the PilT and PilU of *A. baylyi* behave largely like the PilT and PilU of *V. cholerae*.

The *pilU* mutant of *A. baylyi* displays a significant (∼13-fold) reduction in natural transformation (Fig 5), unlike what is observed in *V. cholerae* (Fig 1A). This may be because DNA uptake in *A. baylyi* requires more forceful retraction or because PilU plays a more important role in pilus retraction in this species. Most importantly, however, we found that PilU facilitates natural transformation (and therefore pilus retraction) in backgrounds where the ATPase activity of PilT is putatively inactivated (by mutation of the invariantly conserved lysine in the Walker A domain), but not in backgrounds where *pilT* is absent (Fig 5 – compare *pilT^K136A^* to *pilT^K136A^*Δ*pilU* and Δ*pilT*). Also, a strain where the ATPase activity of both PilT and PilU were putatively inactivated (*pilT^K136A^ pilU^K137A^*) resulted in the loss of transformation equivalent to a retraction deficient background (e.g. Δ*pilTU*) or a strain lacking competence pili (Δ*comP*). Importantly, strains lacking *pilT* (i.e. Δ*pilT* and Δ*pilTU*) could only be complemented via ectopic expression of *pilT* but not *pilU*, while strains that contained an ATPase defective allele of *pilT* (i.e. *pilT^K136A^* Δ*pilU* or *pilT^K136A^ pilU^K137A^*) could be complemented by ectopic expression of either *pilT* or *pilU* (S4 Fig). These data are consistent with what was observed for *V. cholerae* competence pili (Fig 1A and S1 Fig). Additionally, it was very recently shown that PilU from *Pseudomonas aeruginosa* can also promote retraction in a PilT-dependent manner [41]. Together, these results demonstrate that PilU ATPase activity facilitates pilus retraction in a PilT-dependent manner in at least three gammaproteobacteria (*V. cholerae*, *A. baylyi* and *P. aeruginosa*) and suggests that this phenomenon may also extend to other type IVa pilus systems.

**Fig 5.**
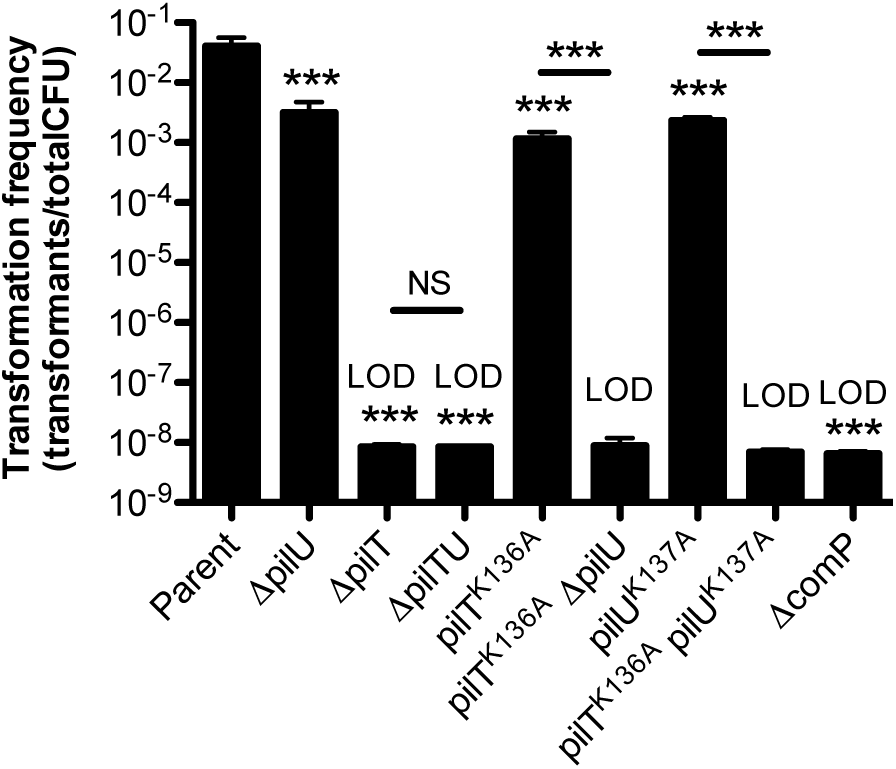
PilU supports pilus retraction in a PilT-dependent manner in *A. baylyi.* Natural transformation assays of the indicated strains incubated with 500ng of transforming DNA for 5 hours. Data are shown as the mean ± SD. Parent, *n* = 5. Δ*comP*, *n* = 6. All other strains, *n* = 4. Asterisk(s) directly above bars denote comparisons to parent strain. All comparisons were made by one-way ANOVA with Tukey’s post test. LOD, limit of detection; NS = not significant; *** = *P* < 0.001.

## Discussion

This study sheds light on the function of PilU as a retraction ATPase of type IVa competence pili in *V. cholerae*. Specifically, our data indicate that PilU can mediate pilus retraction in a PilT-dependent manner likely through its ATPase activity. Our results also show that PilT and PilU form independent homohexamers and that these proteins likely interact with each other and the platform protein PilC. Furthermore, we demonstrate that the role of PilU in supporting pilus retraction is conserved in at least one other type IVa pilus system.

Interestingly, previous work in other species has shown that ATPase mutations in PilT have a greater effect on pilus function compared to the results we present here for both *A. baylyi* and *V. cholerae* competence pili [12, 42]. It is possible that PilT ATPase mutations have a greater effect on retraction depending on the pilus activity in question (for example twitching motility vs DNA uptake). It is also possible that previous reports may have overlooked subtle effects of mutations within the *pilT* gene on the expression of *pilU*, which is generally found downstream in the same operon. Finally, it is also possible that PilU may not be able to compensate for the loss of PilT ATPase activity in some pilus systems.

Our results indicate that PilU can only facilitate pilus retraction in the presence of PilT, regardless of whether that PilT is a functional ATPase. These results are also supported by a recent report from the Blokesch lab, which was published while our manuscript was in revision [41]. We initially tested the hypothesis that this was due to a requirement for PilT to bridge PilU with the platform protein PilC. Our BACTH results, however, indicated that PilU may directly interact with PilC in the absence of PilT (Fig 3 and S2 Fig). While they may interact, it is possible that PilU does not bind and engage the PilC platform in a conformation that would facilitate retraction. Alternatively, it is possible that PilU does bind PilC in a conformation that can facilitate retraction but additional interactions with PilT are required to stimulate PilU ATPase activity. Another possibility is that PilT interacts with other components of the pilus machine that are required for stimulating retraction, and that PilU lacks these interactions. Yet another possibility is that PilU enhances the processivity of the PilT ATPase. Future efforts focused on the interaction network of the retraction ATPases with the pilus machine should shed light on these various hypotheses.

Previous reports have shown that PilU is necessary for twitching motility in *P. aeruginosa* [13], an activity that likely requires forceful retraction [32]. Consistent with this, our results indicate that PilU is necessary for maximal retraction force for the competence pilus in *V. cholerae* (Fig 1G-H). Interestingly, because the Δ*pilU* mutant transforms similarly to the parent strain (Figs 1A and 2B), the forceful retraction mediated by PilU does not seem to be necessary for the function of the *V. cholerae* competence pilus in DNA uptake under the conditions tested. However, forceful PilU-dependent retraction may be necessary for DNA uptake when tDNA is absorbed onto surfaces and/or in complex environments like biofilms.

Beyond altering the force of retraction, many mutants in PilT and PilU also reduced the speed of pilus retraction relative to the parent strain (i.e. Δ*pilU* = ∼1.3-fold, *pilT^K136A^* = ∼4.3-fold, *pilT^K136A^ pilU^L199C^* = ∼6.5-fold, *pilU^K134A^*= ∼1.5-fold, *pilT^L201C^* Δ*pilU* = ∼1.7-fold) (Figs 1B and 2A). This revealed that altering the rates of these motor ATPases results in a corresponding reduction in the rate of pilus retraction [43]. Additionally, despite this reduction in the speed of pilus retraction, these mutants still transformed at levels that were comparable to the parent strain (Figs 1A and 2B). Thus, these results suggest that the absolute speed of competence pilus retraction is not critical for DNA uptake during natural transformation. On the other hand, all mutants that lack a functional retraction ATPase (Δ*pilT*, Δ*pilT*Δ*pilU*, *pilT^K136A^*Δ*pilU*, *pilT^E204A^* Δ*pilU*, *pilT^K136A/E204A^*Δ*pilU*) showed a ∼2-log reduction in the frequency of pilus retraction (Fig 1C). This revealed that functional retraction ATPases are crucial for generating high numbers of retraction events, which may increase the likelihood of retracting a DNA-bound pilus. Further experiments will focus on defining the parameters that are essential for DNA uptake.

Mutants that completely inactivated the retraction machinery (e.g. Δ*pilT*, Δ*pilT* Δ*pilU*, *pilT^K136A^*Δ*pilU*, *pilT^K136A^ pilU^K134A^*) resulted in a hyperpiliated phenotype, lower transformation frequency and a marked reduction in the frequency of pilus retraction (Figs 1A, 1C-E, 2A and 2C). These strains, however, did display rare pilus retraction, which occurred at rates that were markedly slower than the parent (>40-fold), and which was independent of PilT and PilU (Figs 1B and 2A). Recent work in *Neisseria* indicates that PilTU-independent retraction is not unique to the *V. cholerae* type IVa competence pilus [34]. The mechanism underlying PilTU-independent retraction remains unclear. One possibility is that an extension ATPase for another type IV pilus system promotes this retraction. We have tested this in *V. cholerae* by mutating the extension ATPases of the two other type IV pilus systems (MSHA and TCP) and found that PilTU-independent retraction of the competence pilus still occurs in this background (S5 Fig). Thus, suggesting that these other ATPases likely do not contribute to PilTU-independent retraction. It remains possible that the competence pilus extension ATPase PilB supports PilTU-independent retraction as suggested for the type IVc tad pili of *Caulobacter crescentus* where the same ATPase bifunctionally promotes both pilus extension and retraction [43]. It is also possible that PilTU-independent pilus retraction is due to an inherent instability of the pilus fiber. Uncovering the mechanism underlying PilTU-independent retraction of the type IVa competence pilus [5] will be the focus of future work.

## Materials and Methods

### Bacterial strains and culture conditions

*Vibrio cholerae* and *A. baylyi* strains were routinely grown in LB Miller broth and on LB Miller agar supplemented with erythromycin (10 μg/mL), kanamycin (50 µg/mL), and/or carbenicillin (50 µg/mL) when appropriate.

### Construction of mutant strains

All *V. cholerae* strains used throughout this study are derivatives of the El Tor isolate E7946. All *A. baylyi* strains used were derivatives of strain ADP1 [44]. All mutant strains were constructed via MuGENT and natural transformation using transforming DNA products made by splicing-by-overlap extension (SOE) PCR exactly as previously described [45–48]. For a detailed list of all mutant strains used throughout this study see Table S1. For a detailed list of all primers used to construct mutant strains see Table S2.

### Natural transformation assays

In order to induce competence in *V. cholerae* strains, the master competence regulator TfoX was overexpressed using an IPTG-inducible or constitutive P_tac_ promoter and the cells were genetically locked in a state of high cell density via deletion of *luxO* [46, 49–53]. Chitin-independent transformation assays were performed exactly as previously described [47]. Briefly, strains were grown overnight rolling at 30 °C with 100 µM IPTG then ∼10^8^ colony forming units (CFU) were subcultured into 3mL of LB + 100 µM IPTG + 20 mM MgCl_2_ + 10 mM CaCl_2_ and grown to late log. Next, ∼10^8^ CFU of this culture was diluted into instant ocean medium (7 g/L; Aquarium Systems) supplemented with 100 µM IPTG and 500ng of transforming DNA was added to each reaction and incubated statically at 30 °C overnight. The tDNA targeted VC1807, a frame-shifted transposase, for deletion as previously described [45]. Negative control reactions where no tDNA was added were also performed for each strain. After incubation with tDNA, reactions were outgrown by adding 1mL of LB to each reaction and shaking (250 rpm) at 37 °C for ∼2 hours. Reactions were then plated for quantitative culture onto medium selecting for transformants (LB + 10 μg/mL Erythromycin) or onto plain LB for total viable counts. The transformation frequency is defined as the number of transformants divided by the total viable counts. For reactions where no transformants were obtained, a limit of detection was calculated and plotted.

In order to sensitize natural transformation assays to differences in the efficiency of DNA uptake, reactions were incubated with transforming DNA for shorter periods of time. These assays were prepared exactly as described above, except that 10 units of DNase I (NEB) was added to reactions 7 minutes after incubation with 500ng of transforming DNA. Reactions were then incubated statically at 30°C overnight, outgrown, and plated exactly as described above.

For *A. baylyi*, transformations were performed exactly as previously described [54]. Briefly, strains were grown overnight in LB medium. Then, ∼10^8^ cells were diluted into fresh LB medium, and tDNA was added (∼100 ng). Reactions were incubated at 30°C with agitation for 5 h and then plated for quantitative culture as described above to determine the transformation frequency.

### Pilin labelling, imaging and quantification

In order to label pili for observation with epifluorescence microscopy, strains were grown to the late-log phase exactly as described above for natural transformation assays. Then, ∼10^8^ CFU were spun down at 18,000 x g for 1 min and resuspended in instant ocean medium. Cells were then incubated with 25 μg/mL AF488-mal for 15 minutes statically at room temperature in the dark. Cells were washed twice using 100μL of instant ocean medium and resuspended to a final concentration of ∼10^7^ CFU. Next, 2μL of these labeled cells were placed under an 0.2% gelzan pad (made in instant ocean medium) on a coverslip and imaged on an inverted Nikon Ti-2 microscope with a Plan Apo ×60 objective, a green fluorescent protein filter cube, a Hamamatsu ORCAFlash 4.0 camera and Nikon NIS Elements imaging software. In order to measure retraction events, labelled cells were imaged by time-lapse microscopy in which a phase-contrast (to image cell bodies) and fluorescent (to image labeled pili) image were taken every second for 1-2 minutes or every 10 seconds for 10-15 minutes. Rates were calculated manually using measurement tools in the NIS Elements analysis software. Only pili that were longer than 0.3 μm and completed retracting within the time-lapse window were analyzed.

In order to calculate the number of pili per cell, static images of cell bodies and labeled pili were captured using phase-contrast and epifluorescence microscopy, respectively. Images from 3 independent biological replicates were sectioned into areas containing ∼100-200 cells and the total number of cells and the number of pili displayed on each cell was manually determined. Representative images of the piliation state of strains were gathered and the lookup tables for each phase or fluorescent image were adjusted to the same range.

In order to calculate the retraction frequency, time-lapse videos of cell bodies and labeled pili were captured using phase-contrast and epifluorescence microscopy, respectively. Time-lapses from 3 independent biological replicates were sectioned into areas containing ∼300-1000 cells. The number of cells that exhibited a pilus retraction event and the total number of cells in the field was manually counted. This analysis was constrained to measuring pili that are greater than 0.1 μM because shorter pili cannot be resolved by our pilus labeling approach.

To assess aggregation and pelleting of cultures, strains were grown to late log under the conditions described above. Cultures were then allowed to sit statically at room temperature for 30-60 mins. Images of cultures were then taken against a white background.

### Protein expression and purification

His tagged (6xHis) versions of PilT, PilT^K136A^, PilU and PilU^K134A^ were cloned into a pHis expression vector and verified by DNA sequencing (Eurofins). Vectors were transformed into *E. coli* BL21 DE3 for expression and purification.

For purification of His-PilU and His-PilU^K134A^, cells were grown in a 1L flask at 37°C with aeration to an OD_600_ = 0.6. The culture was then induced by adding IPTG (1 mM) and grown to a final OD_600_ = 4.0 at 30°C with aeration. Cells were harvested by centrifugation at 8,000 x g for 15 min at room temperature and then resuspended in Buffer A [100 mM Tris pH 8.5, 300 mM KCl, 10% glycerol, 20 mM Imidazole] and stored at −80°C. Pellets were thawed on ice and then lysozyme (1 mg/mL) and DNase I (2 mg/L) were added before lysing via sonication with a probe tip sonicator. The soluble fraction was clarified by centrifugation at 12,000 x g for 1 hour at 4°C. The supernatant was loaded onto a nickel-charged HisTrap HP column (1mL; GE) using an ÄKTA FPLC at room temperature. The column was then washed with Buffer A and the protein was eluted with a gradient of Buffer B [100 mM Tris pH 8.5, 300 mM KCl, 10% glycerol, 500 mM imidazole] and fractions were collected on ice. Elution fractions were separated on 15% SDS-PAGE gels and stained with Coomassie Brilliant Blue R-250 (BioRad). Peak fractions were pooled and dialyzed (Fisher, MWCO 12 kDa) into 500 mL Buffer C [100 mM Tris pH 8.5, 300 mM KCl, 10% glycerol] overnight, on a stir plate at 4°C. Then a second round of dialysis was performed using 500 mL Buffer C for 4-6 hours, on a stir plate at 4°C. Following dialysis, the protein concentration was determined using a Bradford Assay and aliquots were stored at −80C.

For purification of His-PilT and His-PilT^K136A^, cells were grown in a 1L flask at 37°C with aeration to an OD_600_ = 0.6. The culture was then induced by adding IPTG (1 mM) and grown overnight at 22°C with aeration. Cells were harvested by centrifugation at 8,000 x g for 15min at room temperature and then resuspended in Buffer A [25 mM Tris pH 8.5, 300 mM KCl, 10% glycerol, 1 mM EDTA, 1 mM MgCl_2_, 5 mM *β*-mercaptoethanol, 20 mM Imidazole] and stored at −80°C. Pellets were thawed on ice and lysozyme (1 mg/mL), DNase I (2 mg/L) and PMSF (1 mM) were added before lysing via sonication with a probe tip sonicator. The soluble fraction was clarified by centrifugation at 12,000 x g for 1 hour at 4°C. The supernatant was loaded onto a nickel-charged HisTrap HP column (1 mL; GE) using an ÄKTA FPLC at room temperature. The column was then washed with Buffer A and the protein was eluted with a gradient of Buffer B [25 mM Tris pH 8.5, 300 mM KCl, 10% glycerol, 1 mM EDTA, 1 mM MgCl_2_, 5 mM *β*-mercaptoethanol, 500 mM Imidazole] and fractions were collected on ice. Elution fractions were separated on 15% SDS-PAGE gels and stained with Coomassie Brilliant Blue R-250 (BioRad). Peak fractions were pooled and dialyzed (Fisher, MWCO 12 kDa) into 500 mL Buffer C [25 mM Tris pH 8.5, 50 mM KCl, 10% glycerol, 1 mM EDTA, 1 mM MgCl_2_, 0.5mM TCEP] overnight, on a stir plate at 4°C. The next day, the eluate was concentrated with an Amicon conical filter (MWCO 30 kDa) at 4,000 x g for 15 min at 4°C and resuspended to the original volume in Buffer C twice. The protein concentration was determined using a Bradford Assay and then aliquots of protein were snap-frozen with liquid nitrogen and stored at −80C.

### Bacterial Adenylate Cyclase Two-hybrid (BACTH) Assays

The Bacterial Adenylate Cyclase Two-Hybrid (BACTH) system was employed to study protein interactions. Gene inserts were amplified by PCR and cloned into pUT18C (Carb^R^) and pKT25 (Kan^R^) vectors to generate N-terminal fusions to the T18 and T25 fragments of adenylate cyclase, respectively. Miniprepped vectors (Qiagen plasmid miniprep kit) were then cotransformed into *E. coli* BTH101 and outgrown with 500 µL LB for 1 hour at 37°C with shaking. Transformations were isolation streak plated on LB plates containing kanamycin 50 µg/mL and carbenicillin 100 µg/mL to select for transformants that had received both plasmids. Strains were then grown statically at 30°C from frozen stocks overnight in LB supplemented with kanamycin 50 µg/mL and carbenicillin 100 µg/mL and 3.5µL of each strain was then spotted onto LB plates containing kanamycin 50 µg/mL, carbenicillin 100µg/mL, 0.5 mM IPTG and 40 µg/mL X-gal and incubated at 30°C for 48 hours.

### BACTH Miller Assays

For BACTH Miller assays strains were prepared by growing the BTH101 strains with the indicated pUT18C and pKT25 vectors statically overnight at 30°C in LB supplemented with kanamycin 50 µg/mL and carbenicillin 100 µg/mL. Then 5 µL of this overnight culture was subcultured into 3mL of LB supplemented with kanamycin 50 µg/mL, carbenicillin 100µg/mL and 0.5 mM IPTG and grown rolling at 30°C overnight. Cultures were then pelleted and resuspended in 1 mL of Z buffer [60 mM Na_2_HPO_4_-7H_2_O, 40 mM NaH_2_PO_4_-H_2_O, 10 mM KCl, 2 mM MgSO_4_-7H_2_O] to an OD_600_ = 0.5. 50µL of 1% SDS was added to each reaction and vortexed. 100µL of chloroform was added to each reaction, vortexed again and then centrifuged for 1 min at 18,000 x g. The aqueous fraction was then mixed with ONPG (2.5 mg/mL) in a 96-well plate and the OD_420_ and OD_550_ were measured every 5 minutes for 10 hours. Miller units were then calculated at the time (in mins) when the A_420_ reached 0.3 or at 10 hours using the formula: [1000*(A_420_-(1.75*A_550_))/ (mins*0.2 mL*A_600_)].

### Western Blotting

Strains were grown to mid-log phase in LB with 100 µM IPTG if applicable and then resuspended to an OD_600_ = 100 in TBS containing 1 mM PMSF. Cells were then lysed using a FastPrep-24 (MP Bio). Lysates were centrifuged at 16,000 x g for 10 min at 4°C, and the supernatant was then mixed 1:1 with 2X SDS sample buffer [220 mM Tris pH 6.8, 25% glycerol, 1.8% SDS, 0.02% Bromophenol Blue, 5% *β*-mercaptoethanol] and then 2uL of each sample was separated on a 15% SDS PAGE gel. Proteins were transferred to a PVDF membrane and incubated with one of the following primary antibodies: α-FLAG polyclonal rabbit (Sigma) or α-RpoA monoclonal mouse (Biolegend). Then, blots were washed and incubated with a α-rabbit or α-mouse secondary antibodies conjugated to IRdye 800CW. Blots were then imaged using an Odyssey classic LI-COR system.

### Negative-stain electron microscopy

Negative stain specimens were prepared by applying 4 µL of protein solution (0.1 mg/mL + 1 mM AMP-PnP) onto a glow-discharged 300-mesh copper grid coated with continuous carbon film (EMS). After 30 s the protein solution was blotted with a piece of filter paper. The grid was washed with 4 µL of milli-Q water and stained with 4 µL of stain solution (1% (w/v) uranyl acetate, 0.5% (w/v) trehalose). The excess stain solution was removed by filter paper and the grid was left to air dry. Due to the availability of the instrument, we used two different transmission electron microscopes for data collection. For PilU, the data were collected on a 300-kV JEM-3200FS TEM equipped with FEG source and 8k x 8k Direct Electron DE-64 camera. A total of 32 micrograph images were taken at a nominal magnification of 60,000x, which is equivalent to 1.56 Å per pixel. For PilT, the data were collected on a 120-kV JEM-1400plus TEM equipped with LaB_6_ source and a Gatan Oneview camera. A total of 59 images were taken at a nominal magnification of 60,000x and the pixel size is 1.94 Å. To avoid potential beam induced damage, minimum dose system was employed during data collection. Note that PilU data were collected on a direct electron detector; therefore, each image contains 50 frames and the frame alignment was performed using unblur program with exposure filter applied.

Particle picking, defocus estimation and phase flipped particles were all done using EMAN2 software package [55]. These particles were then imported into RELION (v3.0.7) for 2D classification, Initial model building, 3D refinement, and 3D classification [56]. From 2D classification results, we observed clear hexameric density organization [7, 57]. No symmetry was applied for subsequent 3D refinement and 3D classification. The final 3D reconstructions were computed from 45204 particles to 16.6 Å for PilU and 17472 particles to 20.8 Å for PilT. The resolution estimation was calculated using gold-standard method at Fourier shell correlation of 0.143 as implemented in RELION. The 3D reconstructions were rendered using UCSF Chimera where the handedness of PilU was chosen to be consistent with PilT [58]. The negative stain 3D reconstruction of PilU and PilT have been deposited into EMDB (ID: EMD-20757, EMD-20758 respectively).

### Micropillar retraction assay

To ensure that all measurements recorded were due to type IV competence pili, all the experiments were performed with *V*. *cholerae* strains lacking all external appendages other than type IV competence pili as well as exopolysaccharide production (i.e. mutants lacking flagella, MSHA pili, TCP pili and *Vibrio* polysaccharide (VPS) production). Strains were grown overnight in LB either directly from frozen stocks or from a single colony from an LB agar plate. Of the overnight liquid culture, 50 μl was subcultured into 3 ml LB + 100 μM IPTG + 20 mM MgCl_2_ + 10 mM CaCl_2_ and allowed to grow at 30°C for 4h. Then, 100 μl of this culture was centrifuged for 5 min at 20,000 x *g* and the bacteria were resuspended in instant ocean medium. At different times, 23 μl of the resuspension were added to micropillars in an observation chamber as previously reported [15]. Briefly, silica molds were inverted on activated coverslips with polyacrylamide gels in between. The result is an array of flexible micropillars in a hexagonal array of 3 μm × 3 μm.Once the bacteria were in contact with the micropillars, 10-Hz movies of the top of the pillars were recorded. The motion of the tips of the pillars was tracked using a cross-correlation algorithm in ImageJ. The amplitude and speeds of the pillars’ motions were then analyzed using MATLAB. Finally, we calibrated the pillars’ stiffness constant using optical tweezers as previously described [33]. The pillars used in this study have a stiffness constant of 17±4pNμm−1.

### Statistics

Statistical differences were assessed by one-way ANOVA tests followed by a multiple comparisons post test (either Tukey’s or Dunnett’s) using GraphPad Prism software. For all transformation frequency experiments and frequency of pilus retraction experiments, statistical analyses were performed on log transformed data. All means and statistical comparisons can be found in S3 and S4 Tables.

## Acknowledgements

We would like to thank CK Ellison for helpful discussions. We would like to acknowledge JD Newman and CE Dann III for their advice during protein purification. We thank JC van Kessel for providing strains associated with BACTH assays. We would like to thank L Khosla and V Deopersaud for technical assistance with micropillar experiments. We also thank JA Lorah for assistance with statistical analyses.

## Supporting Information

**S1 Fig.**
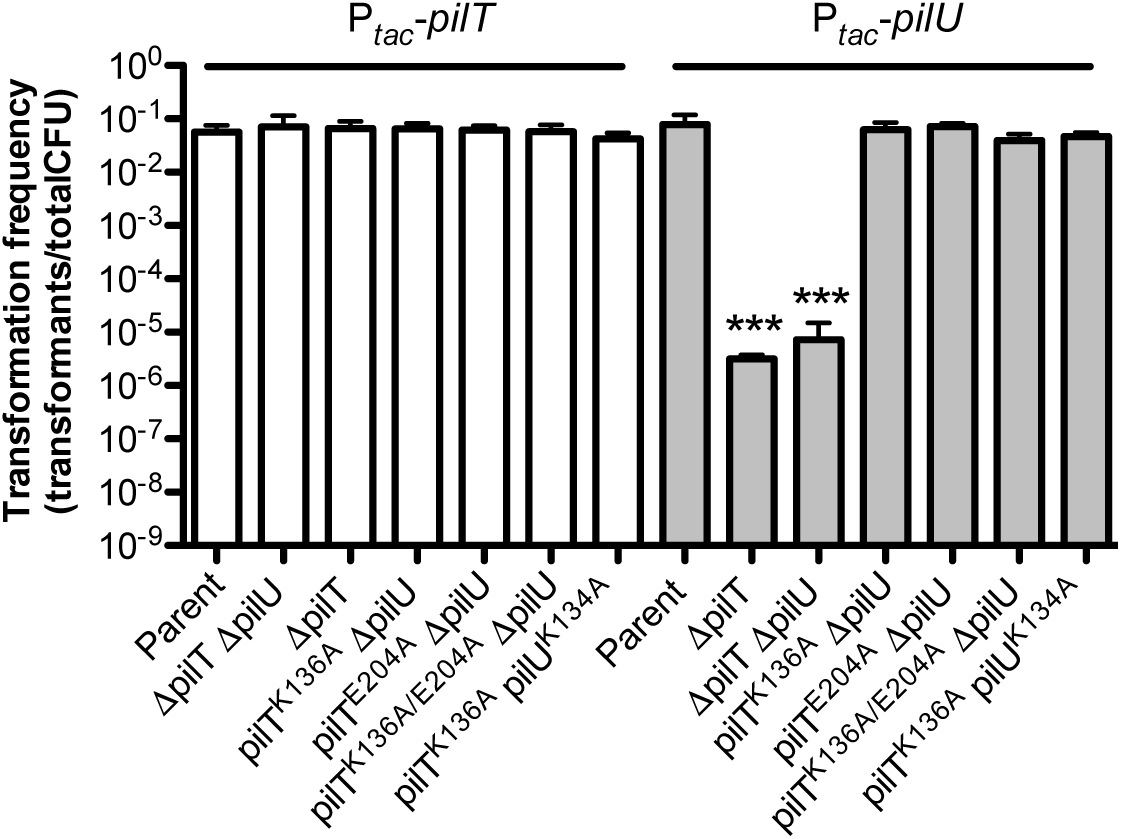
Complementation analysis of *V. cholerae* mutant strains via ectopic expression of *pilT* or *pilU*. Natural transformation assays of the indicated strains. All strains contain either a chromosomally integrated IPTG-inducible P*_tac_*-*pilT* (white bars) or P*_tac_*-*pilU* (gray bars) construct as indicated. Data show that ectopic overexpression of PilT and PilU does not affect the transformation frequency of the parent strain. Ectopic overexpression of PilT rescued the transformation of all strains that showed a substantial reduction in natural transformation in Fig 1A and Fig 2B (i.e. Δ*pilT*, Δ*pilT* Δ*pilU*, *pilT^K136A^* Δ*pilU*, *pilT^E204A^* Δ*pilU*, *pilT^E204A/K136A^* Δ*pilU*, and *pilT^K136A^ pilU^K134A^*). Ectopic overexpression of PilU rescued all strains except for strains that lacked a copy of *pilT* (i.e. Δ*pilT*, Δ*pilT* Δ*pilU*). All strains were grown with 100 µM IPTG to ectopically induce PilT or PilU. Data are shown as the mean ± SD. Parent, *n* = 8. All other strains, *n* = 4. Asterisk(s) directly above bars denote comparisons to the appropriate parent strain. Comparisons were made by grouping data into the two families indicated by the white and grey bars. A one-way ANOVA was performed for each family with a Dunnet’s post to compare each experimental sample to the parent strain. Comparisons were not statistically significantly different unless otherwise noted. *** = *P* < 0.001.

**S2 Fig.**
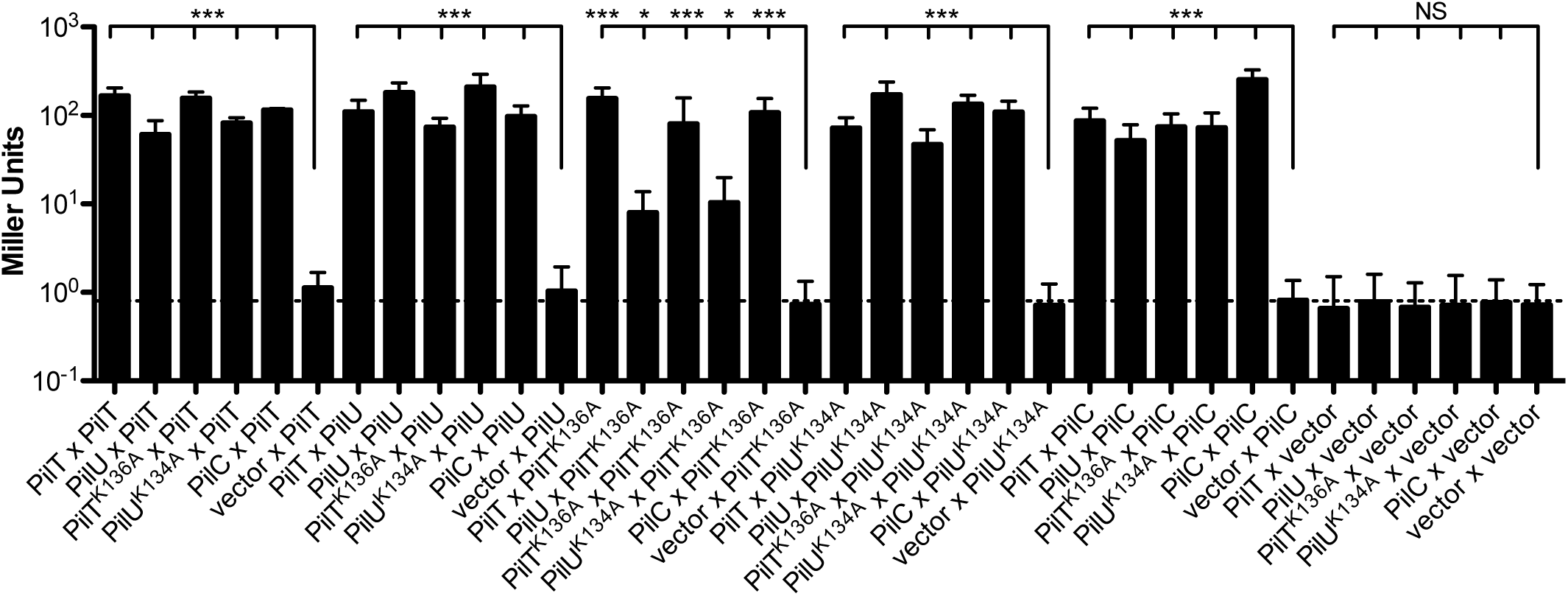
PilT and PilU interact with each other and the pilus platform protein PilC. BACTH Miller assays showing all data from Fig 3B with additional negative controls for every T25- or T18-fusion. Data are shown as the mean ± SD from three independent experiments. The dotted line represents the β-galactosidase activity of the empty vector negative control (T18-empty x T25-empty). Comparisons were made by grouping data into appropriate families indicated by the black lines above the bar graph. A one-way ANOVA was performed on each family with a Dunnet’s post test to compare each experimental group to the appropriate negative control (indicated by the longer black line). A single set of asterisks directly above the group line denotes the statistical results for all comparisons in that family unless otherwise noted. NS, not significant; * = *P* < 0.05; *** = *P* < 0.001.

**S3 Fig.**
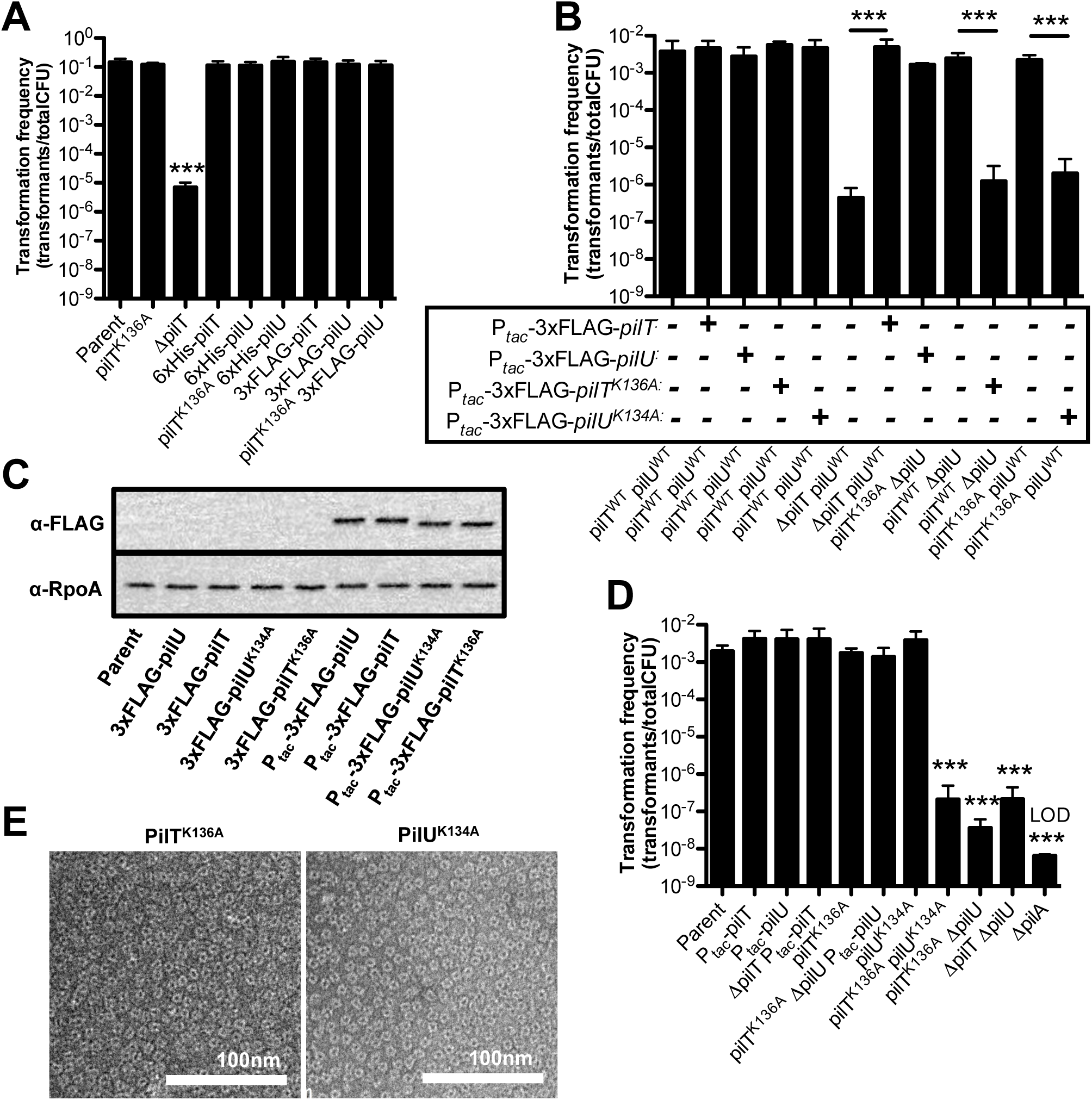
PilT and PilU form homomeric complexes to facilitate pilus retraction. **(A)** Natural transformation assays showing that 6xHis and 3xFLAG N-terminal fusions to PilT and PilU are functional. The indicated strains were incubated with 500 ng of transforming DNA overnight. Parent, *n* = 8. All others, *n* = 4. **(B)** Natural transformation assays showing that overexpression of 3xFLAG tagged PilT/PilU/PilT^K136A^/PilU^K134A^ behave the same as non-tagged proteins when ectopically overexpressed in the indicated backgrounds. Strains were incubated with 500 ng of transforming DNA and after 7 minutes, DNAse I was added to prevent additional DNA uptake. Parent, *n* = 5. *pilT^K136A^*, *n* = 5. All others, *n* = 4. **(C)** Western blot of the indicated strains to detect FLAG tagged-proteins and RpoA as a loading control. Strains containing an IPTG regulated P*_tac_* constuct were grown in the presence of 100 µM IPTG. This blot indicates that ectopic induction of 3xFLAG tagged PilT/PilU/PilT^K136A^/ PilU^K134A^ results in robust overexpression of proteins above native levels. Data is representative of three independent experiments. **(D)** Natural transformation assays where the indicated strains were incubated with 500 ng of transforming DNA for 7 minutes prior to the addition of DNAse I to prevent additional DNA uptake. These data indicate that ectopic overexpression of PilT or PilU can rescue strains with a substantial reduction in transformation frequency. All strains with P*_tac_* constructs were grown with 100 µM IPTG to overexpress PilT or PilU. All bar graphs are shown as the mean ± SD. Asterisk(s) directly above bars denote comparisons to parent strain. All comparisons were made by one-way ANOVA followed with Tukey’s post test. LOD, limit of detection; *** = *P* < 0.001. **(E)** Representative negative stain transmission electron micrographs showing that purified ^6X^His-PilT^K136A^ and ^6X^His-PilU^K134A^ form hexamers *in vitro.* Scale bar, 100 nm.

**S4 Fig.**
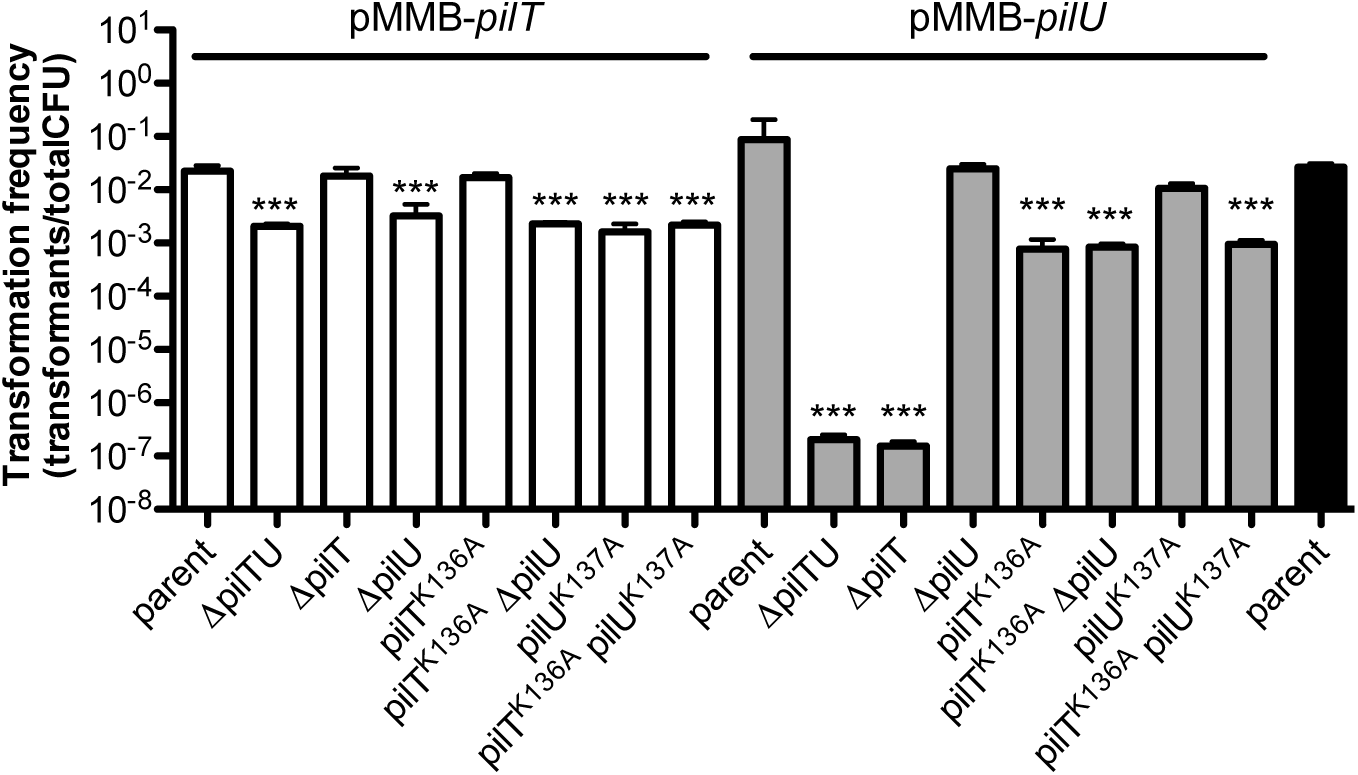
Complementation analysis of *A. baylyi* mutant strains via ectopic expression of *pilT* or *pilU*. Natural transformation assays of the indicated *A. baylyi* strains. Strains harbored pMMB-*pilT* (white bars), pMMB-*pilU* (gray bars), or no vector (black bar). Data indicate that ectopic overexpression of PilT and PilU do not affect the transformation frequency of the parent strain. Ectopic overexpression of PilT rescued the transformation of all strains that showed a significant reduction in natural transformation in Fig 5 (i.e. Δ*pilT*, Δ*pilT* Δ*pilU*, *pilT^K136A^* Δ*pilU*, and *pilT^K136A^ pilU^K137A^*). Ectopic overexpression of PilU rescued all strains except for strains that lacked a copy of pilT (i.e. Δ*pilT* and Δ*pilT* Δ*pilU*). All strains were grown with 100 µM IPTG to induce expression of *pilT* or *pilU*. Data are shown as the mean ± SD and are from three independent experiments. Asterisk(s) directly above bars denote comparisons to the appropriate parent strain. Comparisons were made by grouping data into the two families indicated by the white and grey bars. A one-way ANOVA was performed for each family followed by Dunnet’s post test to compare each experimental group to the parent strain. Comparisons were not statistically significantly different from the relevant parent unless otherwise noted. *** = *P* < 0.001.

**S5 Fig.**
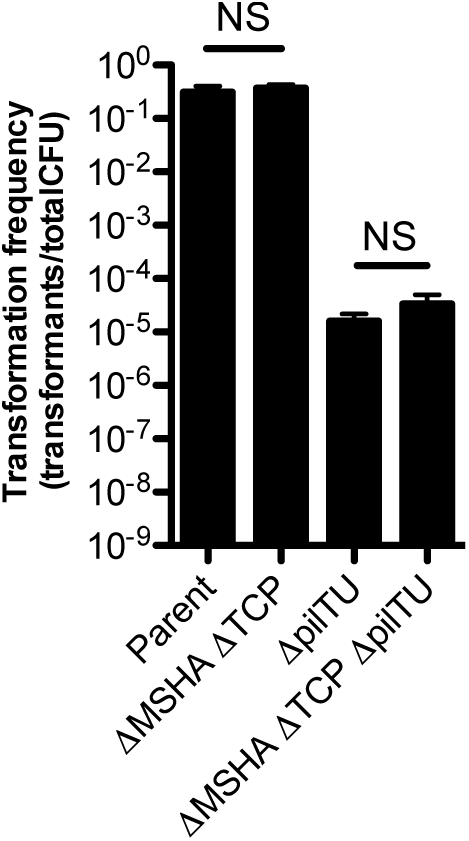
The components of the *V. cholerae* MSHA or TCP type IV pilus systems do not contribute to PilTU-independent retraction of the competence pilus. Natural transformation assays of the indicated strains. Reactions were incubated with 500 ng of transforming DNA overnight. ΔMSHA and ΔTCP represent deletions of the entire locus for both pilus systems, which includes the extension ATPase associated with each. Data are shown as the mean ± SD and are from four independent biological replicates. All comparisons were made by one-way ANOVA with Tukey’s post test. NS, not significant.

**S1 Table.**
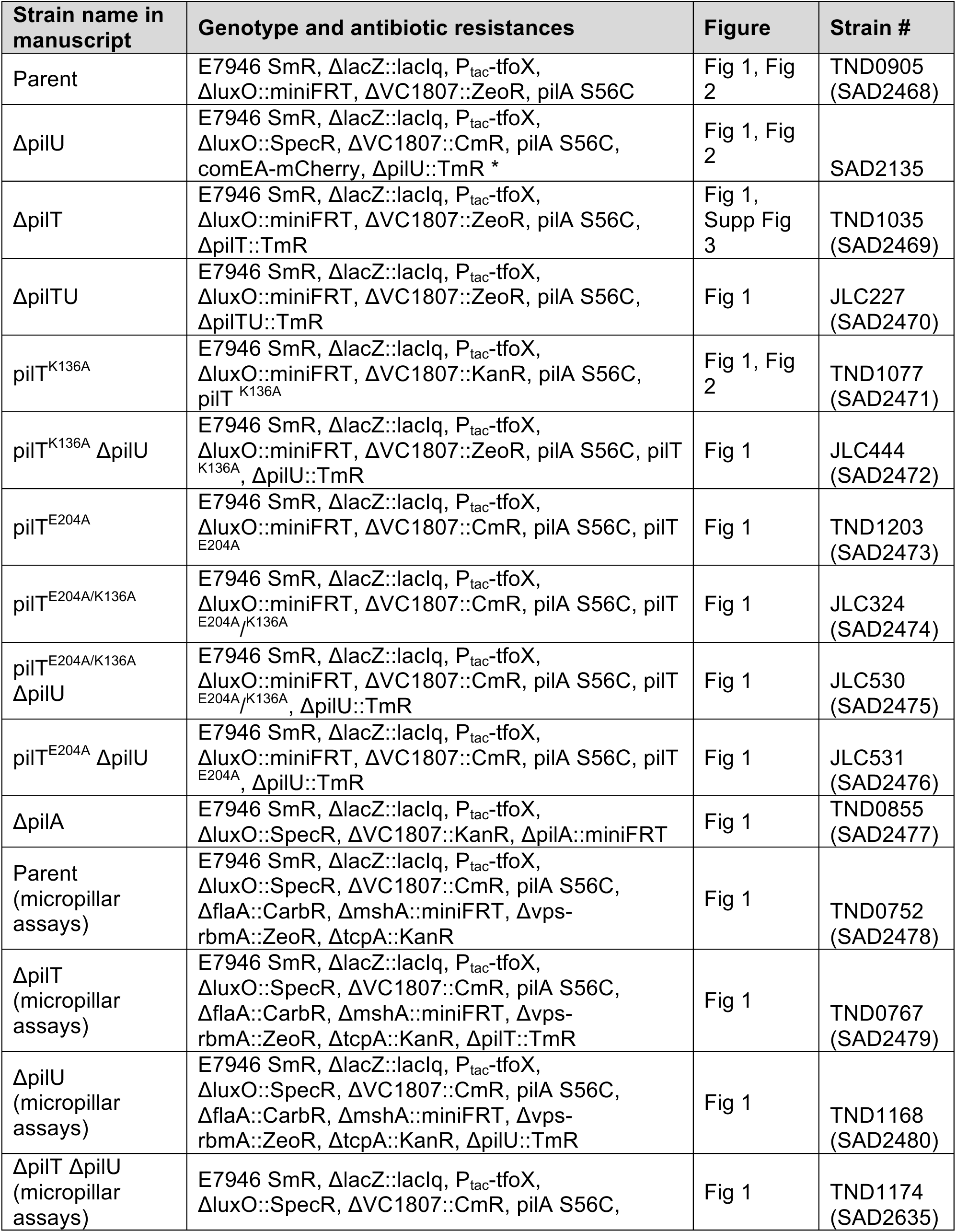
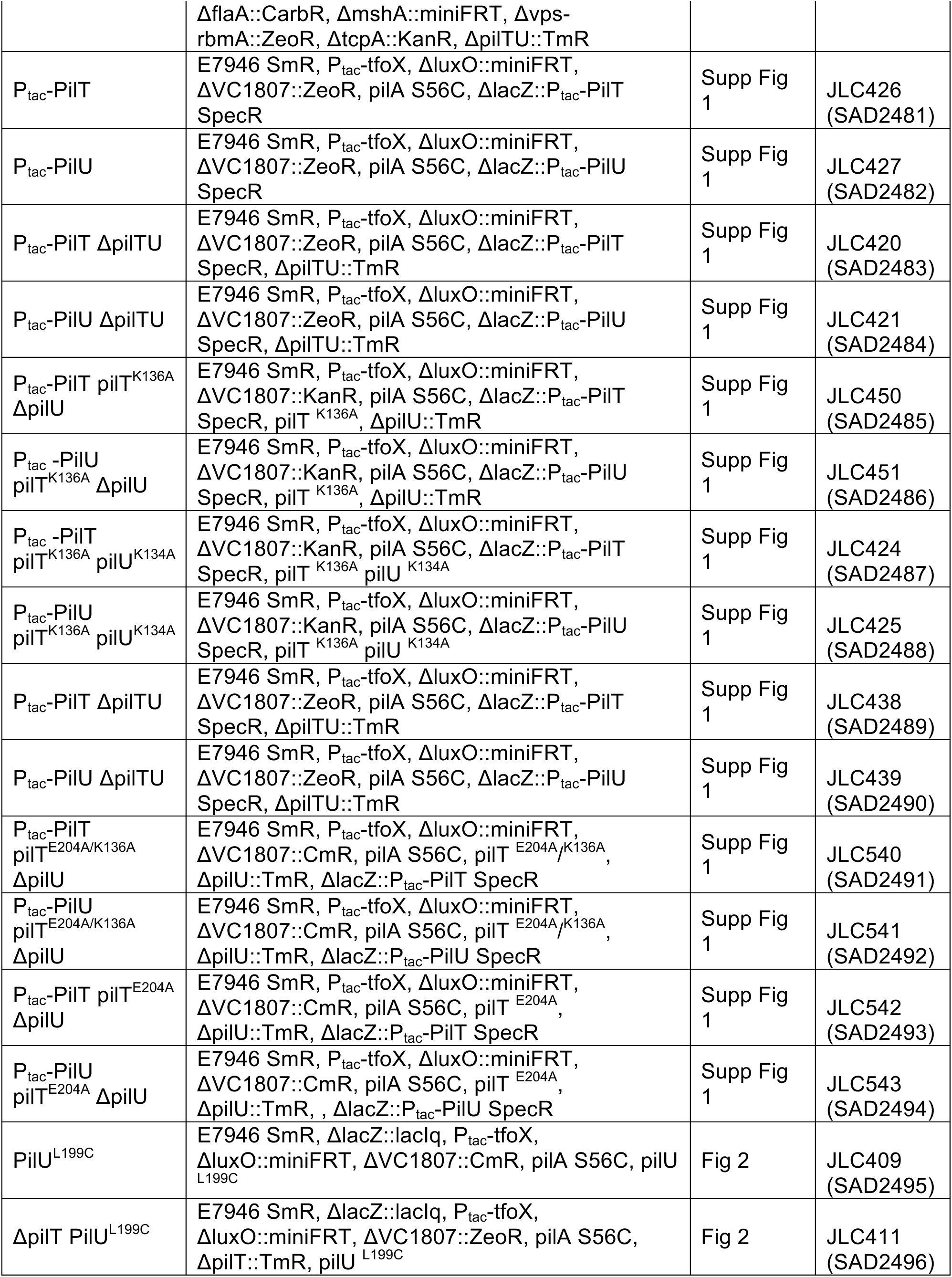
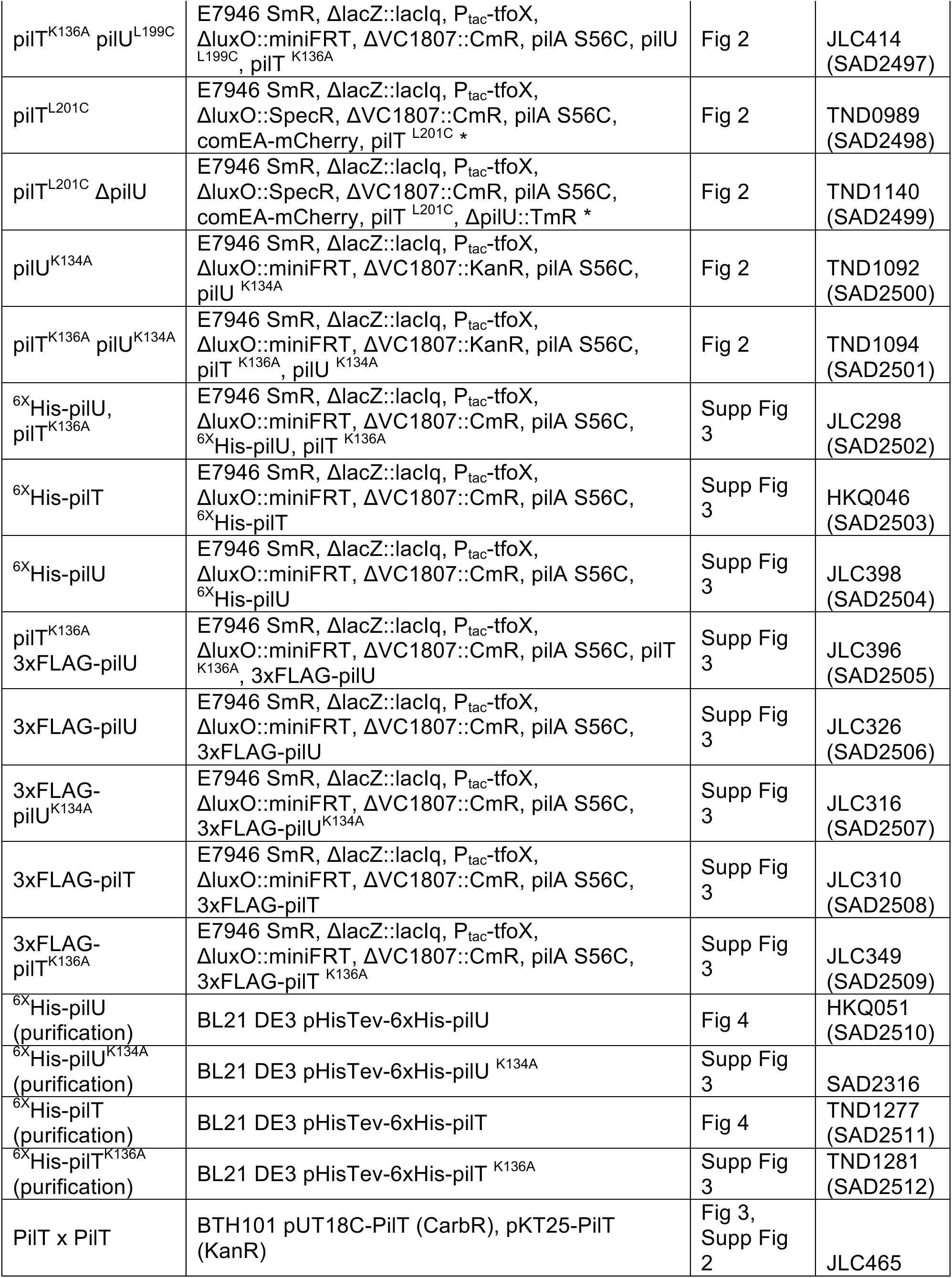
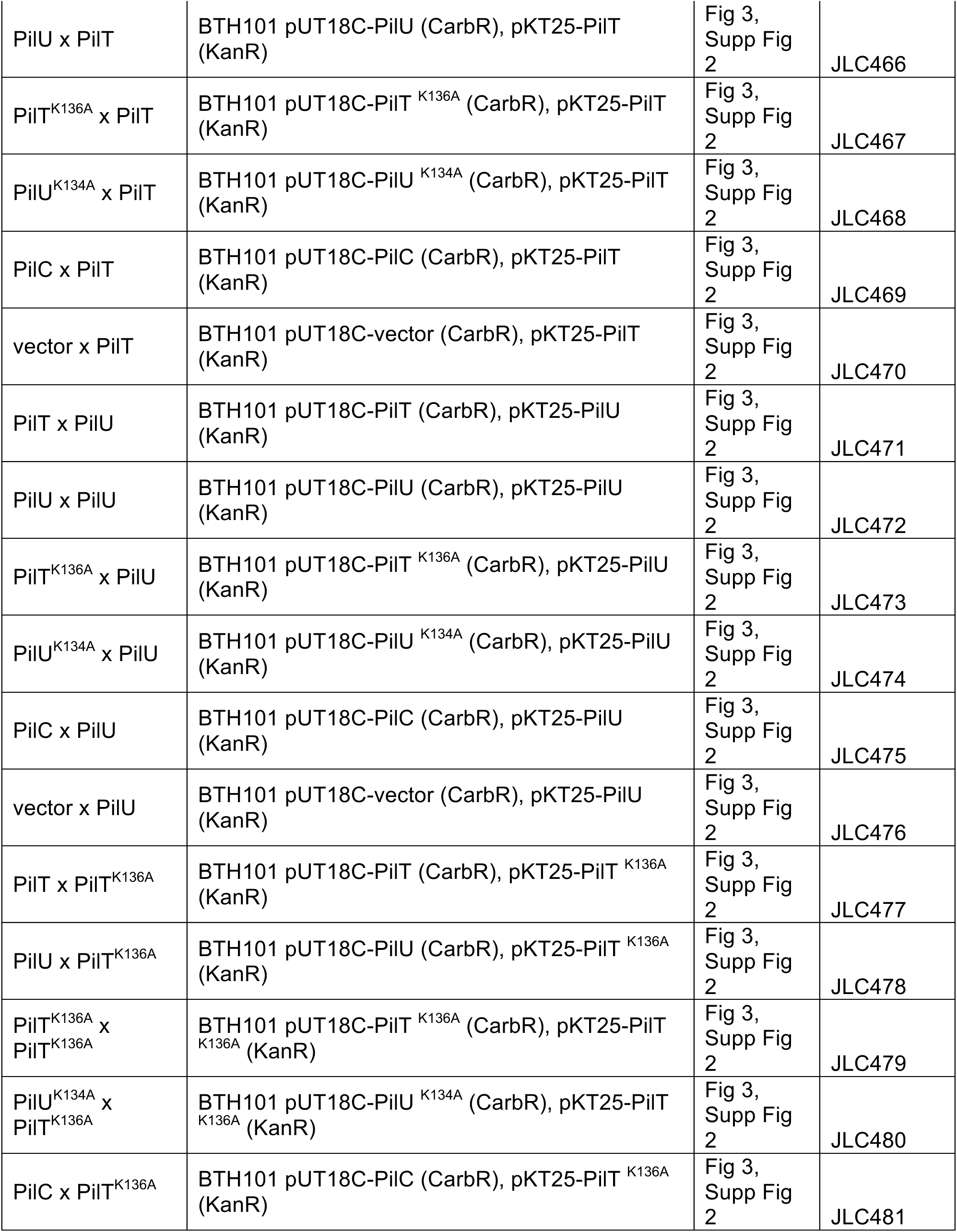
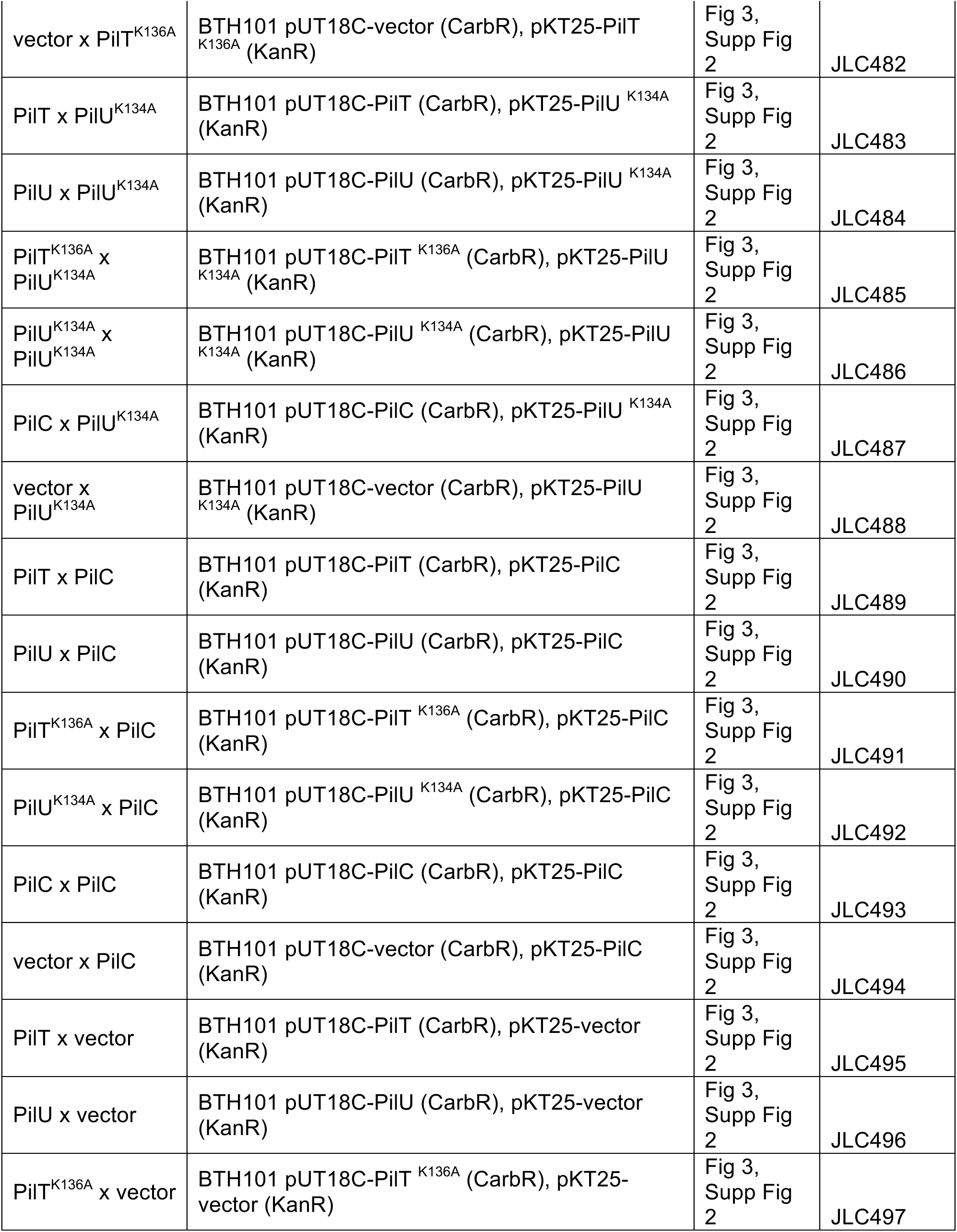
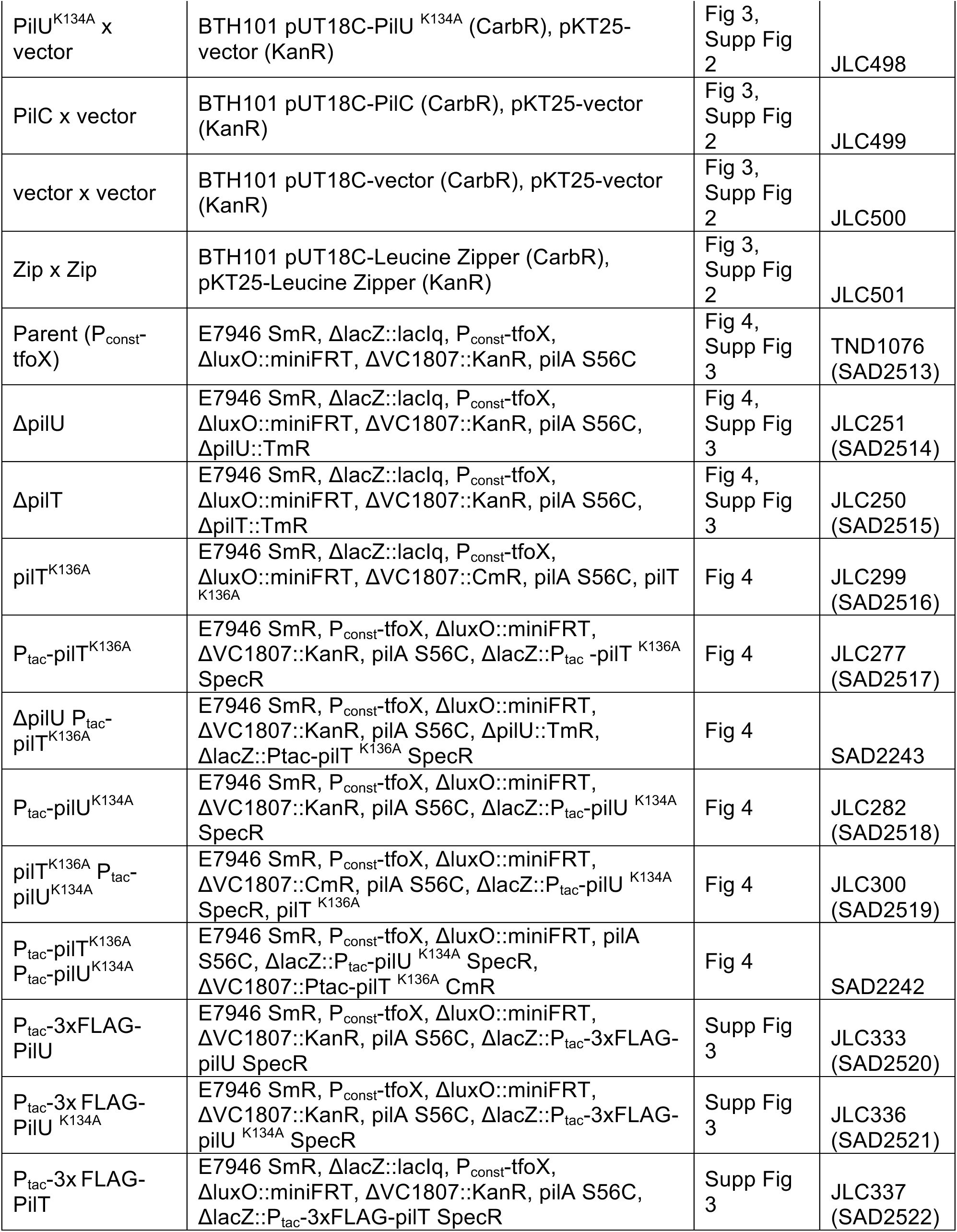
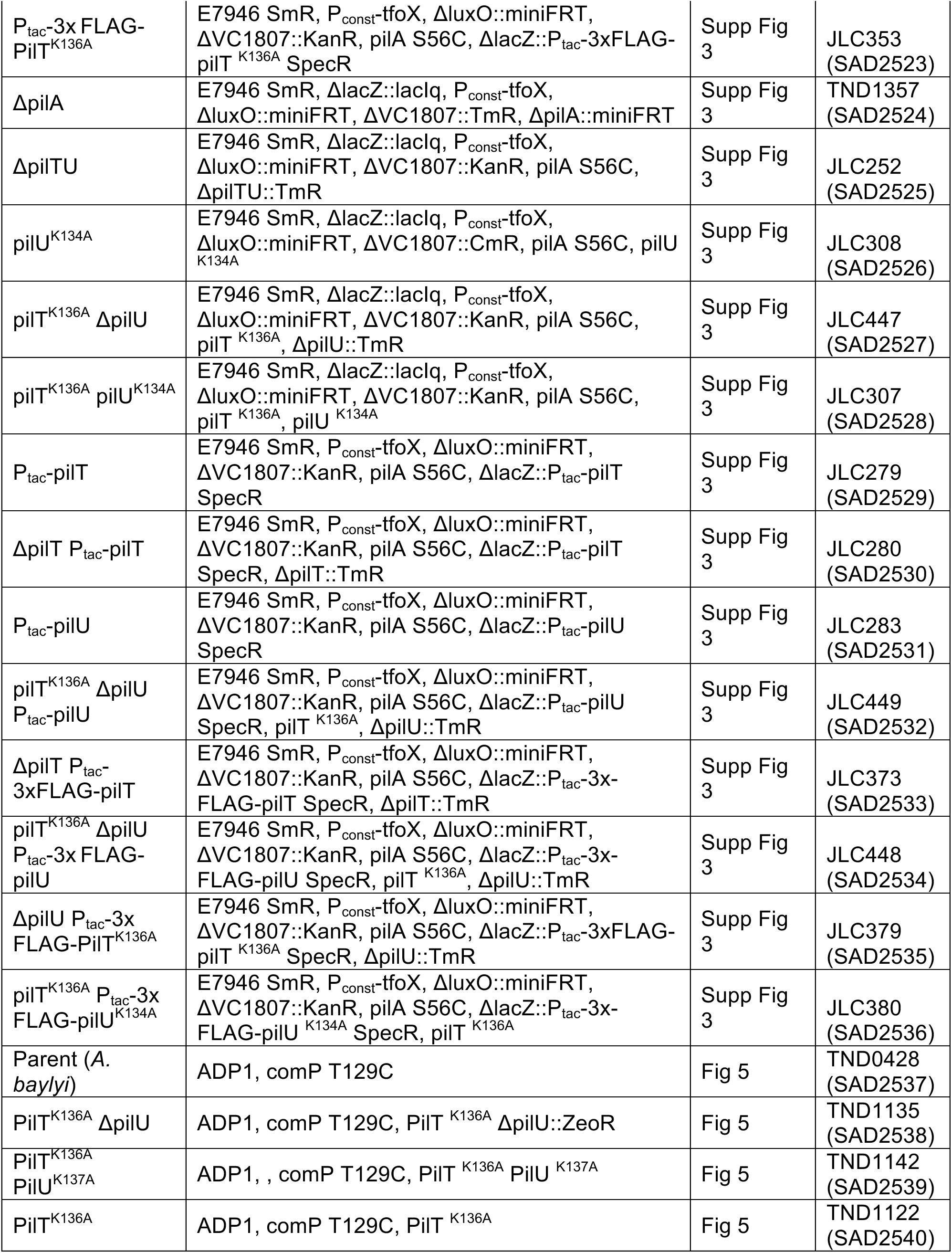
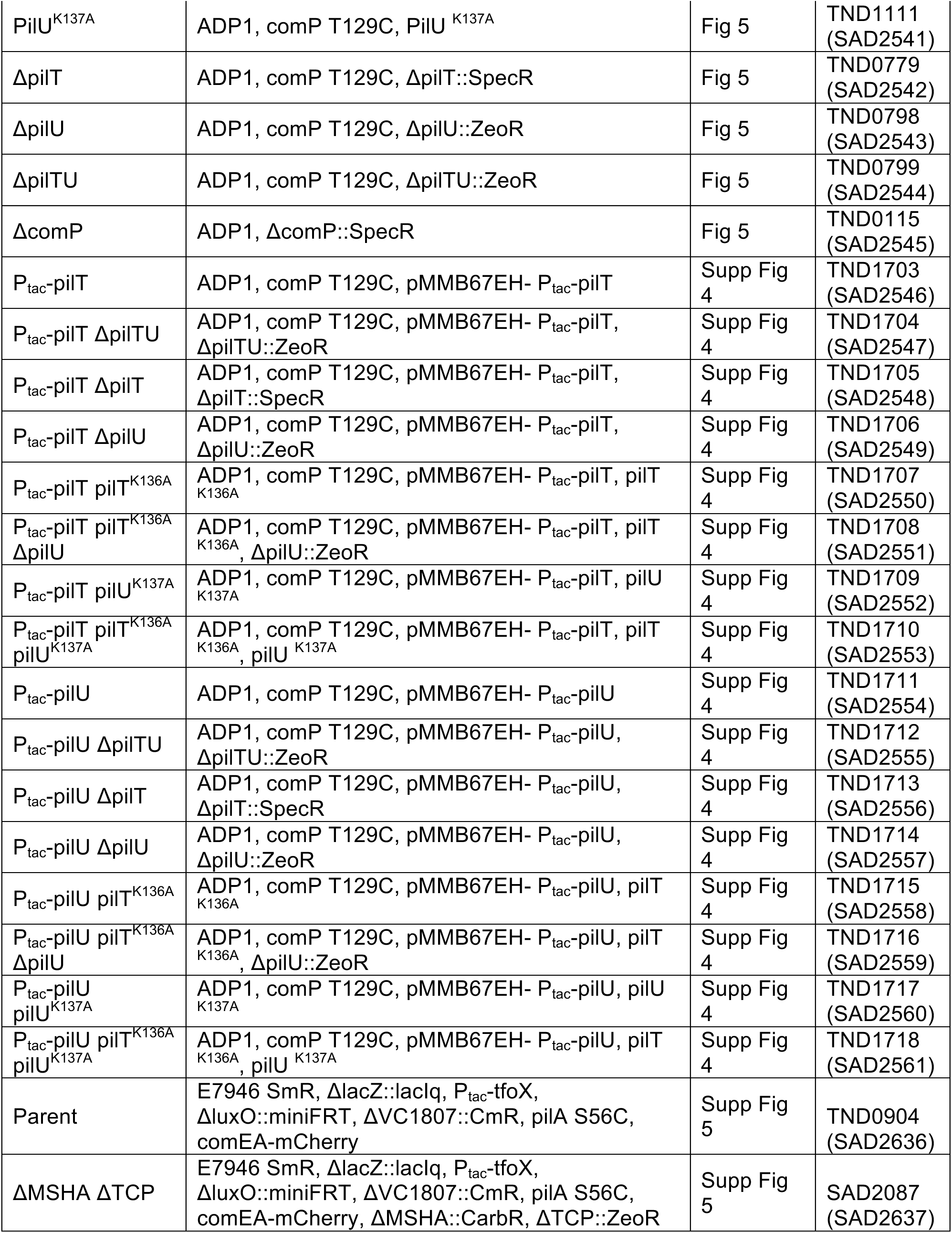
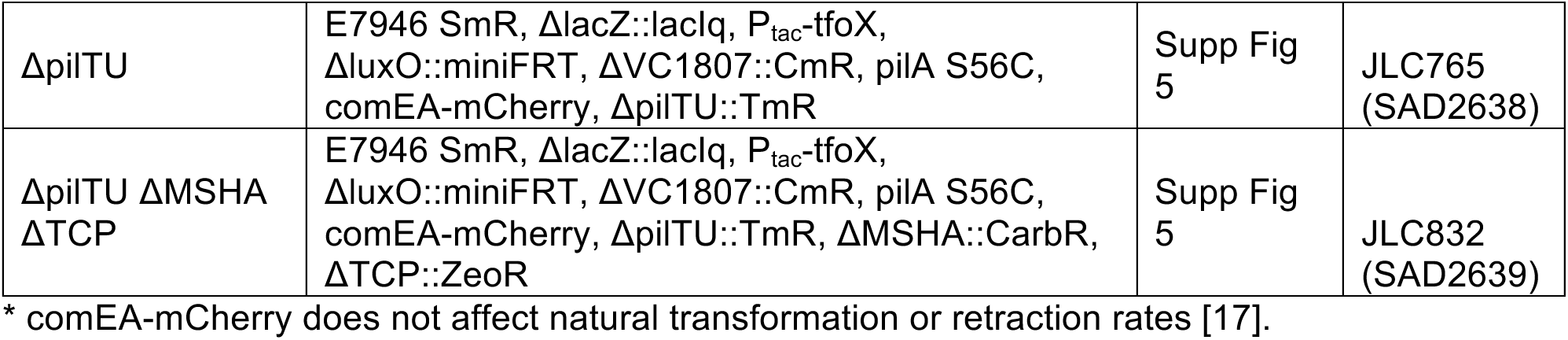
Strains used in this study.

**S2 Table.**
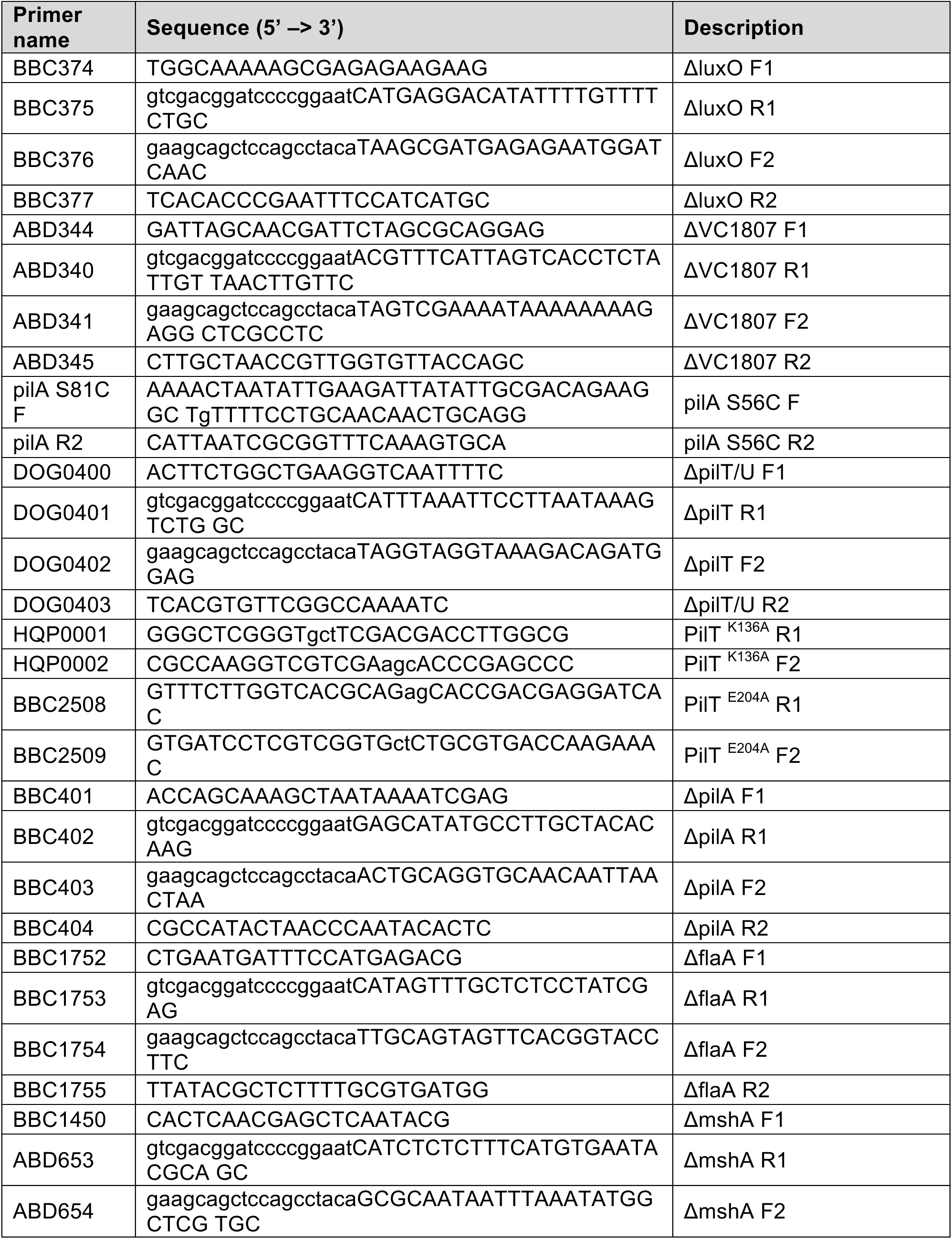
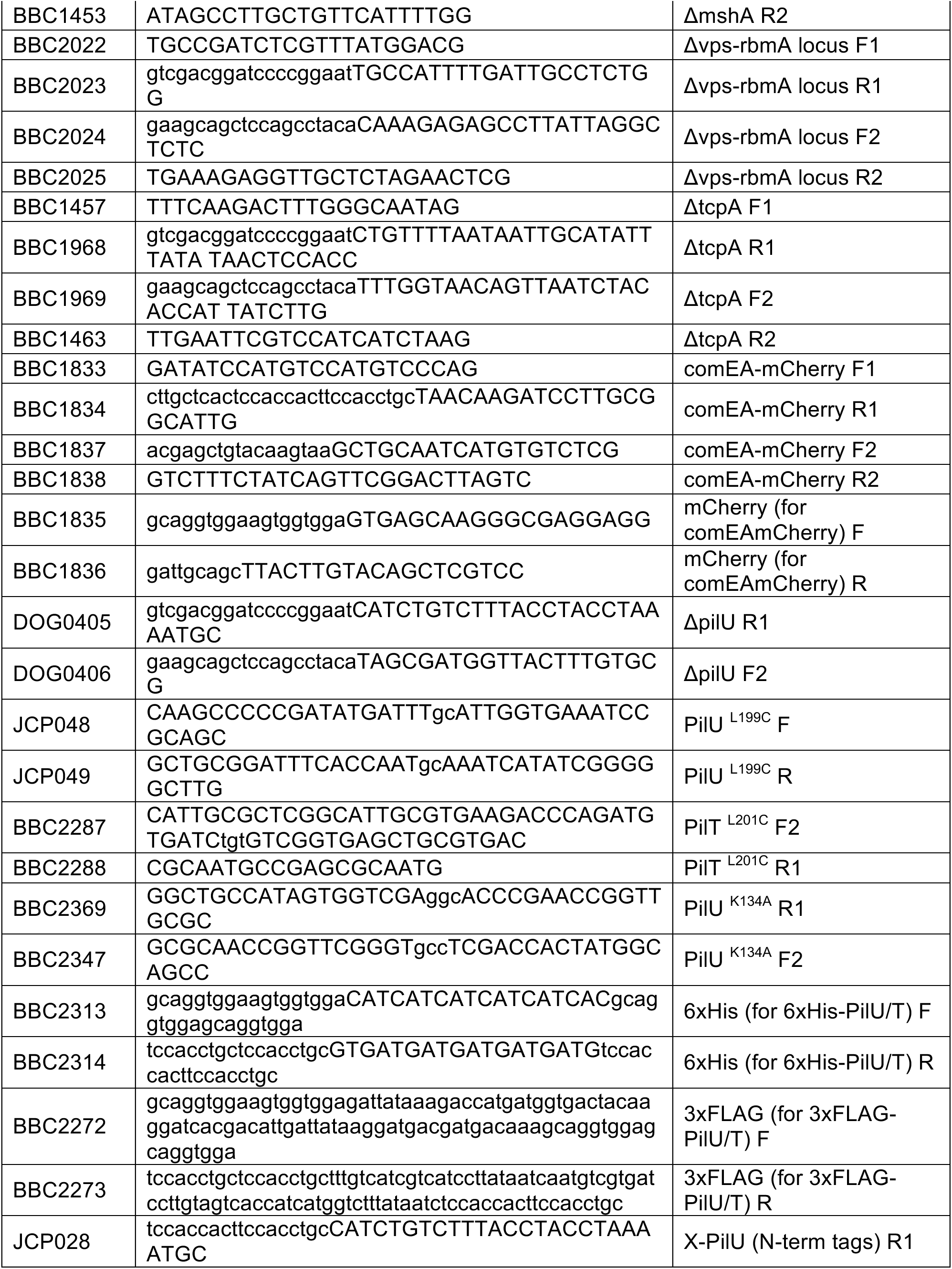
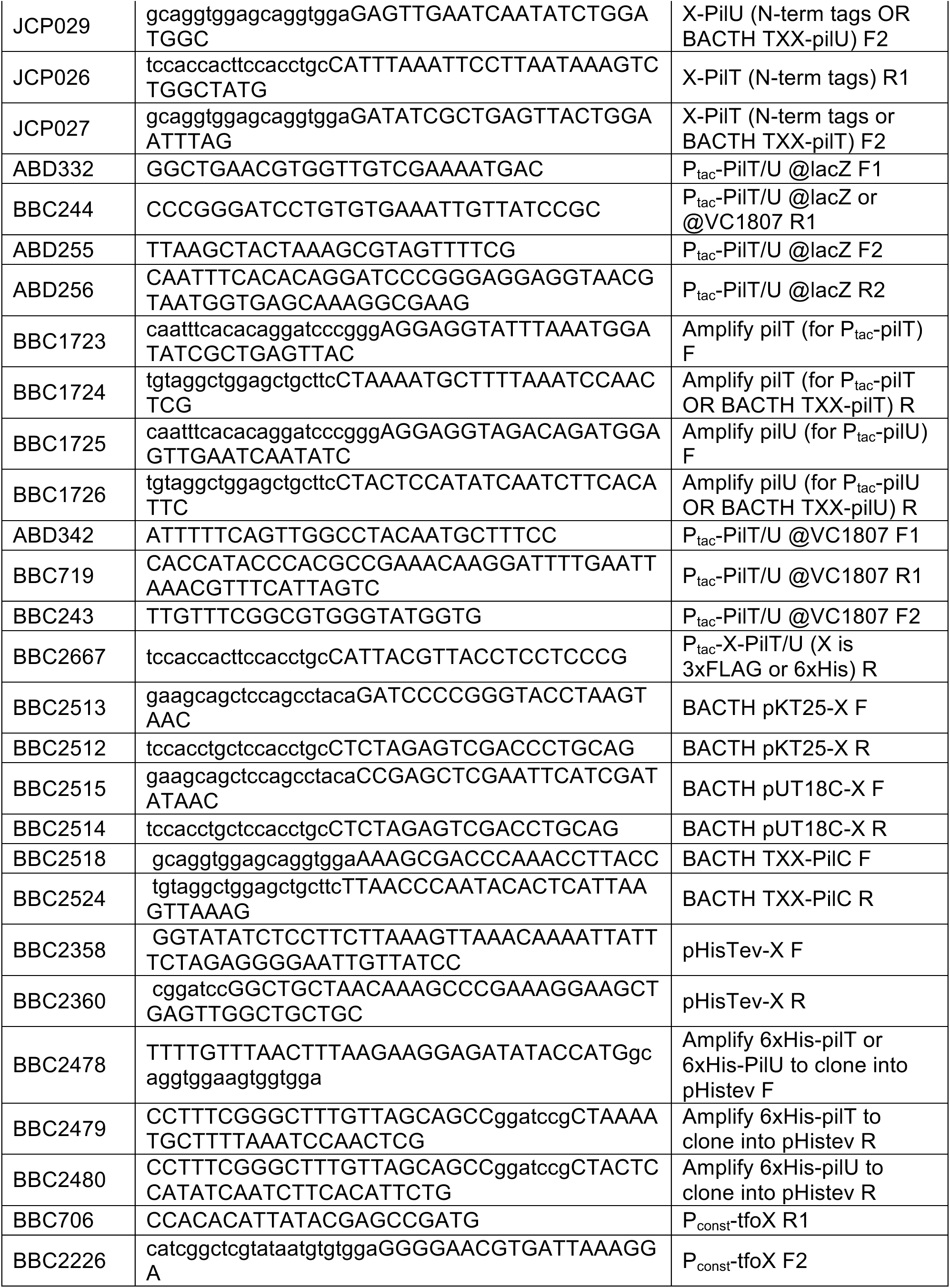
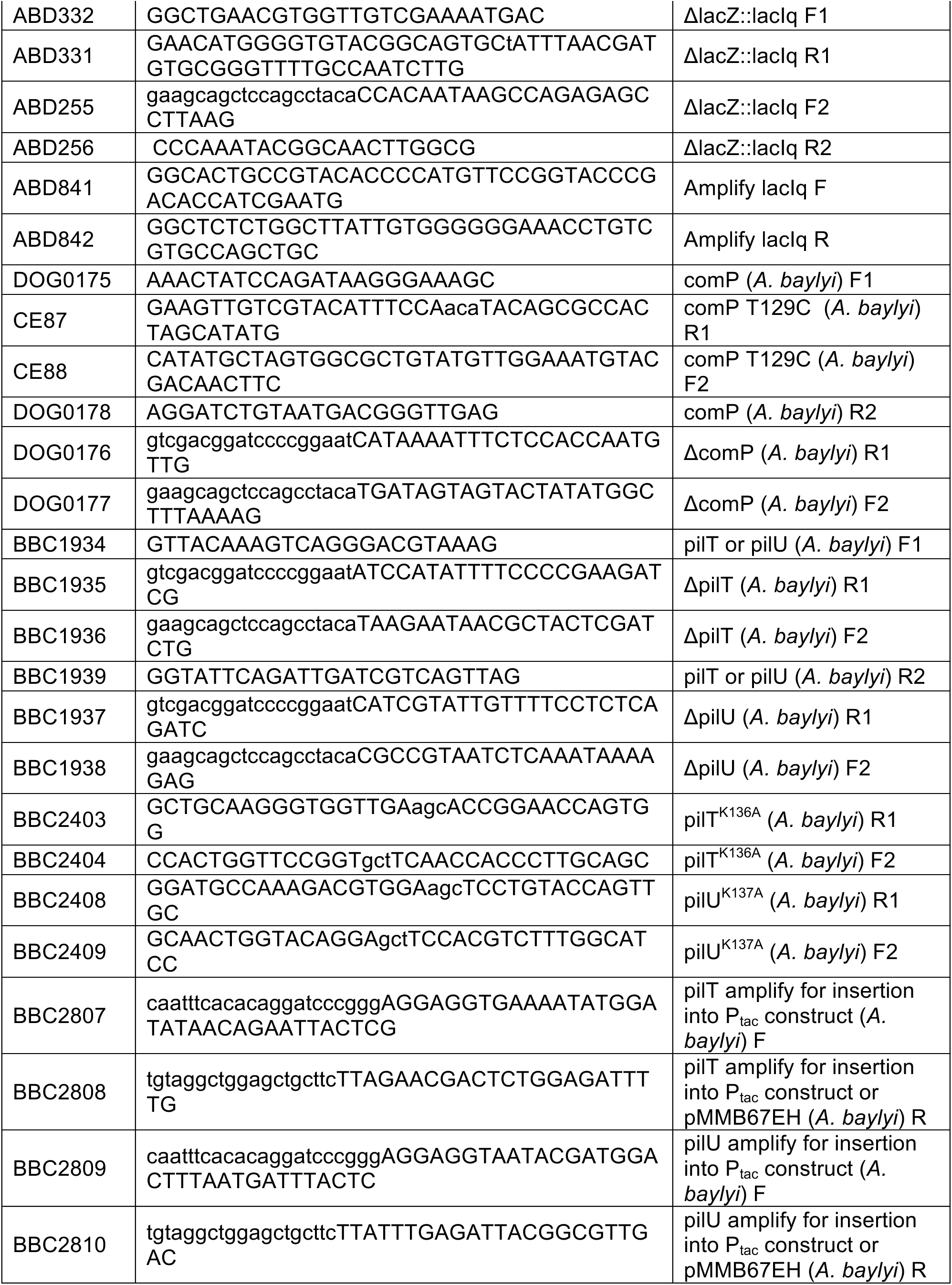
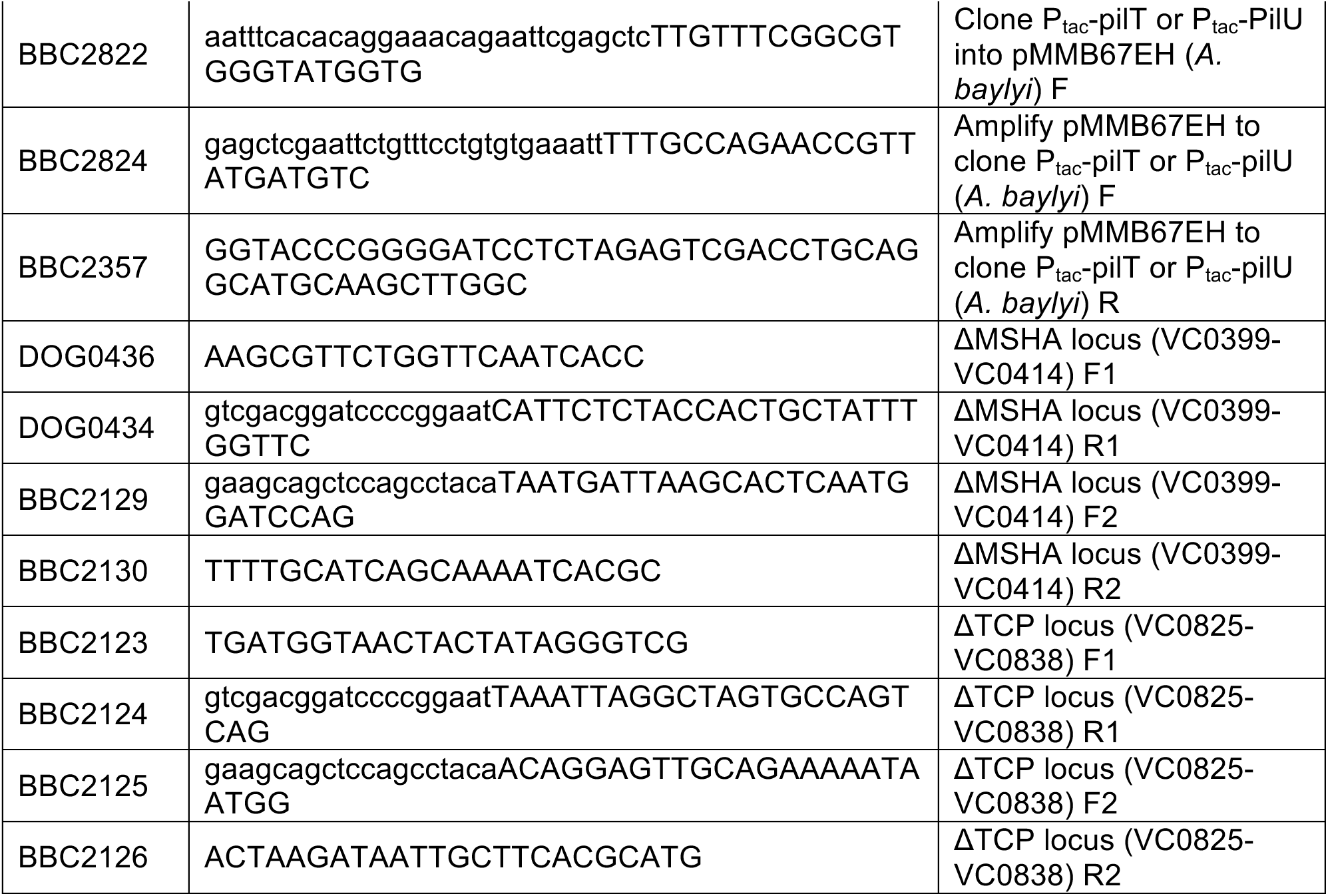
Primers used in this study.

**S3 Table.** Mean values for each data set.

| Figure | Strain Name | Mean | Figure | Strain Name | Mean |
| --- | --- | --- | --- | --- | --- |
| Fig 1A | Parent | 1.08E-01 | S2 Fig | PilT x PilT | 1.67E+02 |
| Fig 1A | ΔpilA | 1.22E-08 | S2 Fig | PilU x PilT | 6.10E+01 |
| Fig 1A | ΔpilU | 5.78E-02 | S2 Fig | PilT<sup>K136A</sup> x PilT | 1.57E+02 |
| Fig 1A | ΔpilT | 3.29E-06 | S2 Fig | PilU<sup>K134A</sup> x PilT | 8.23E+01 |
| Fig 1A | ΔpilTΔpilU | 3.91E-06 | S2 Fig | PilC x PilT | 1.15E+02 |
| Fig 1A | pilT<sup>K136A</sup> | 6.22E-02 | S2 Fig | vector x PilT | 1.13E+00 |
| Fig 1A | pilT<sup>K136A</sup> ΔpilU | 3.89E-06 | S2 Fig | PilT x PilU | 1.10E+02 |
| Fig 1A | pilT<sup>E204A</sup> | 2.55E-02 | S2 Fig | PilU x PilU | 1.81E+02 |
| Fig 1A | pilT<sup>K136A/E204A</sup> | 3.53E-02 | S2 Fig | PilT<sup>K136A</sup> x PilU | 7.37E+01 |
| Fig 1A | pilT<sup>E204A</sup> ΔpilU | 9.36E-06 | S2 Fig | PilU<sup>K134A</sup> x PilU | 2.11E+02 |
| Fig 1A | pilT<sup>K136A/E204A</sup> ΔpilU | 4.66E-06 | S2 Fig | PilC x PilU | 9.71E+01 |
| Fig 1B | Parent | 2.12E-01 | S2 Fig | vector x PilU | 1.04E+00 |
| Fig 1B | ΔpilU | 1.57E-01 | S2 Fig | PilT x PilT<sup>K136A</sup> | 1.56E+02 |
| Fig 1B | ΔpilT | 3.07E-03 | S2 Fig | PilU x PilT<sup>K136A</sup> | 7.99E+00 |
| Fig 1B | ΔpilTΔpilU | 3.85E-03 | S2 Fig | PilT<sup>K136A</sup> x PilT<sup>K136A</sup> | 8.06E+01 |
| Fig 1B | pilT<sup>K136A</sup> | 4.26E-02 | S2 Fig | PilU<sup>K134A</sup> x PilT<sup>K136A</sup> | 1.04E+01 |
| Fig 1B | pilT<sup>K136A</sup> ΔpilU | 2.06E-03 | S2 Fig | PilC x PilT<sup>K136A</sup> | 1.08E+02 |
| Fig 1B | pilT<sup>E204A</sup> | 4.82E-02 | S2 Fig | vector x PilT<sup>K136A</sup> | 7.32E-01 |
| Fig 1B | pilT<sup>E204A/K136A</sup> | 6.12E-02 | S2 Fig | PilT x PilU<sup>K134A</sup> | 7.20E+01 |
| Fig 1B | pilT<sup>E204A</sup> ΔpilU | 3.58E-03 | S2 Fig | PilU x PilU<sup>K134A</sup> | 1.72E+02 |
| Fig 1B | pilT<sup>E204A/K136A</sup> ΔpilU | 4.09E-03 | S2 Fig | PilT<sup>K136A</sup> x PilU<sup>K134A</sup> | 4.69E+01 |
| Fig 1C | Parent | 1.73E+01 | S2 Fig | PilU<sup>K134A</sup> x PilU<sup>K134A</sup> | 1.34E+02 |
| Fig 1C | ΔpilU | 3.16E+01 | S2 Fig | PilC x PilU<sup>K134A</sup> | 1.10E+02 |
| Fig 1C | ΔpilT | 2.93E-02 | S2 Fig | vector x PilU<sup>K134A</sup> | 7.14E-01 |
| Fig 1C | ΔpilTΔpilU | 1.33E-01 | S2 Fig | PilT x PilC | 8.69E+01 |
| Fig 1C | pilT<sup>K136A</sup> | 3.43E+01 | S2 Fig | PilU x PilC | 5.20E+01 |
| Fig 1C | pilT<sup>K136A</sup> ΔpilU | 8.75E-02 | S2 Fig | PilT<sup>K136A</sup> x PilC | 7.43E+01 |
| Fig 1C | pilT<sup>E204A</sup> | 1.41E-01 | S2 Fig | PilU<sup>K134A</sup> x PilC | 7.25E+01 |
| Fig 1C | pilT<sup>E204A/K136A</sup> | 1.77E+01 | S2 Fig | PilC x PilC | 2.55E+02 |
| Fig 1C | pilT<sup>E204A</sup> ΔpilU | 5.65E-02 | S2 Fig | vector x PilC | 8.15E-01 |
| Fig 1C | pilT<sup>E204A/K136A</sup> ΔpilU | 3.86E-02 | S2 Fig | PilT x vector | 6.59E-01 |
| Fig 1F | Parent | 1.05E+01 | S2 Fig | PilU x vector | 7.85E-01 |
| Fig 1F | ΔpilT | 4.93E+00 | S2 Fig | PilT<sup>K136A</sup> x vector | 6.81E-01 |
| Fig 1F | ΔpilU | 5.08E+00 | S2 Fig | PilU<sup>K134A</sup> x vector | 7.14E-01 |
| Fig 1F | ΔpilTΔpilU | 6.08E+00 | S2 Fig | PilC x vector | 7.68E-01 |
| Fig 1H | Parent | 1.93E-01 | S2 Fig | vector x vector | 7.22E-01 |
| Fig 1H | ΔpilT | 1.37E-02 | S3-A Fig | Parent | 1.46E-01 |
| Fig 1H | ΔpilU | 7.23E-02 | S3-A<br>Fig | ΔpilT | 6.94E-06 |
| Fig 1H | ΔpilTΔpilU | 8.19E-03 | S3-A<br>Fig | pilT<sup>K136A</sup>, 6xHis-pilU | 1.53E-01 |
| Fig 2A | Parent | 1.37E-01 | S3-A<br>Fig | 6xHis-pilT | 1.14E-01 |
| Fig 2A | pilU<sup>L199C</sup> | 1.51E-01 | S3-A<br>Fig | 6xHis-pilU | 1.11E-01 |
| Fig 2A | pilT<sup>K136A</sup> pilU<sup>L199C</sup> | 9.81E-02 | S3-A<br>Fig | pilT<sup>K136A</sup>, 3xFLAG-pilU | 1.14E-01 |
| Fig 2A | ΔpilT pilU<sup>L199C</sup> | 6.79E-06 | S3-A<br>Fig | 3xFLAG-pilT | 1.45E-01 |
| Fig 2A | pilT<sup>L201C</sup> | 2.04E-01 | S3-A<br>Fig | 3xFLAG-pilU | 1.23E-01 |
| Fig 2A | pilT<sup>L201C</sup> ΔpilU | 6.47E-02 | S3-A<br>Fig | pilT<sup>K136A</sup> | 1.20E-01 |
| Fig 2A | pilT<sup>K136A</sup> | 1.17E-01 | S3-B<br>Fig | Parent | 3.75E-03 |
| Fig 2A | pilU<sup>K134A</sup> | 6.33E-02 | S3-B<br>Fig | ΔpilT | 4.46E-07 |
| Fig 2A | pilT<sup>K136A</sup> pilU<sup>K134A</sup> | 6.07E-07 | S3-B<br>Fig | Ptac-3xFLAG-pilT | 4.60E-03 |
| Fig 2A | ΔpilU | 8.80E-02 | S3-B<br>Fig | Ptac-3xFLAG-pilT ΔpilT | 4.94E-03 |
| Fig 2B | Parent | 2.12E-01 | S3-B<br>Fig | Ptac-3xFLAG-pilU | 2.78E-03 |
| Fig 2B | ΔpilU | 1.57E-01 | S3-B<br>Fig | Ptac-3xFLAG-pilU, pilT<sup>K136A</sup><br>ΔpilU | 1.66E-03 |
| Fig 2B | pilT<sup>K136A</sup> | 4.26E-02 | S3-B<br>Fig | Ptac-3xFLAG-pilT<sup>K136A</sup> | 5.66E-03 |
| Fig 2B | pilU<sup>K134A</sup> | 1.37E-01 | S3-B<br>Fig | Ptac-3xFLAG-pilU<sup>K134A</sup> | 4.71E-03 |
| Fig 2B | pilT<sup>K136A</sup> pilU<sup>K134A</sup> | 4.77E-03 | S3-B<br>Fig | Ptac-3xFLAG-pilT<sup>K136A</sup>,<br>ΔpilU | 1.25E-06 |
| Fig 2B | pilT<sup>K136A</sup> pilU<sup>L199C</sup> | 3.26E-02 | S3-B<br>Fig | P<sub>tac</sub>-3xFLAG-pilU<sup>K134A</sup>,<br>pilT<sup>K136A</sup> | 2.00E-06 |
| Fig 2B | pilU<sup>L199C</sup> | 1.96E-01 | S3-B<br>Fig | ΔpilU | 2.48E-03 |
| Fig 2B | pilT<sup>L201C</sup> | 2.11E-01 | S3-B<br>Fig | pilT<sup>K136A</sup> | 2.24E-03 |
| Fig 2B | pilT<sup>L201C</sup> ΔpilU | 1.28E-01 | S3-D<br>Fig | Parent | 1.97E-03 |
| Fig 2B | ΔpilT pilU<sup>L199C</sup> | 3.91E-03 | S3-D<br>Fig | ΔpilT ΔpilU | 2.15E-07 |
| Fig 4C | Parent | 2.02E-03 | S3-D<br>Fig | pilU<sup>K134A</sup> | 3.90E-03 |
| Fig 4C | ΔpilU | 2.57E-03 | S3-D<br>Fig | pilT<sup>K136A</sup> ΔpilU | 3.62E-08 |
| Fig 4C | ΔpilT | 1.57E-06 | S3-D<br>Fig | pilT<sup>K136A</sup> pilU<sup>K134A</sup> | 2.12E-07 |
| Fig 4C | pilT<sup>K136A</sup> | 3.08E-03 | S3-D<br>Fig | P<sub>tac</sub>-pilT | 4.18E-03 |
| Fig 4C | Ptac-pilT<sup>K136A</sup> | 5.94E-03 | S3-D<br>Fig | ΔpilT Ptac-pilT | 4.08E-03 |
| Fig 4C | Ptac-pilT<sup>K136A</sup> ΔpilU | 2.41E-06 | S3-D Fig | P<sub>tac</sub>-pilU | 4.07E-03 |
| Fig 4C | Ptac-pilU<sup>K134A</sup> | 5.55E-03 | S3-D Fig | pilT<sup>K136A</sup> ΔpilU Ptac-pilU | 1.39E-03 |
| Fig 4C | Ptac-pilU<sup>K134A</sup> pilT<sup>K136A</sup> | 2.46E-05 | S3-D Fig | ΔpilA | 6.49E-09 |
| Fig 4C | Ptac-pilT<sup>K136A</sup> Ptac-pilU<sup>K134A</sup> | 4.82E-06 | S3-D Fig | pilT<sup>K136A</sup> | 1.75E-03 |
| Fig 5 | pilT<sup>K136A</sup> ΔpilU | 8.95E-09 | S4 Fig | Parent pmmB-pilT | 2.23E-02 |
| Fig 5 | pilT<sup>K136A</sup> pilU<sup>K137A</sup> | 7.00E-09 | S4 Fig | ΔpilTU pmmB-pilT | 2.05E-03 |
| Fig 5 | pilT<sup>K136A</sup> | 1.17E-03 | S4 Fig | ΔpilT pmmB-pilT | 1.80E-02 |
| Fig 5 | pilU<sup>K137A</sup> | 2.34E-03 | S4 Fig | ΔpilU pmmB-pilT | 3.24E-03 |
| Fig 5 | Parent | 4.10E-02 | S4 Fig | pilT<sup>K136A</sup> pmmB-pilT | 1.68E-02 |
| Fig 5 | ΔpilT | 8.52E-09 | S4 Fig | pilT<sup>K136A</sup> ΔpilU pmmB-pilT | 2.30E-03 |
| Fig 5 | ΔpilU | 3.17E-03 | S4 Fig | pilU<sup>K137A</sup> pmmB-pilT | 1.63E-03 |
| Fig 5 | ΔpilTU | 8.48E-09 | S4 Fig | pilT<sup>K136A</sup> pilU<sup>K137A</sup> pmmB-pilT | 2.19E-03 |
| Fig 5 | Δcomp | 6.53E-09 | S4 Fig | Parent pmmB-pilU | 8.67E-02 |
| S1 Fig | ΔpilT ΔpilU Ptac-PilT | 7.05E-02 | S4 Fig | ΔpilTU pmmB-pilU | 2.04E-07 |
| S1 Fig | ΔpilT ΔpilU Ptac-PilU | 7.26E-06 | S4 Fig | ΔpilT pmmB-pilU | 1.55E-07 |
| S1 Fig | pilT<sup>K136A</sup> ΔpilU Ptac-PilT | 6.53E-02 | S4 Fig | ΔpilU pmmB-pilU | 2.48E-02 |
| S1 Fig | pilT<sup>K136A</sup> ΔpilU Ptac-PilU | 6.27E-02 | S4 Fig | pilT<sup>K136A</sup> pmmB-pilU | 7.66E-04 |
| S1 Fig | pilT<sup>K136A</sup> pilU<sup>K134A</sup> Ptac-PilT | 4.21E-02 | S4 Fig | pilT<sup>K136A</sup> ΔpilU pmmB-pilU | 8.35E-04 |
| S1 Fig | pilT<sup>K136A</sup> pilU<sup>K134A</sup> Ptac-PilU | 4.65E-02 | S4 Fig | pilU<sup>K137A</sup> pmmB-pilU | 1.07E-02 |
| S1 Fig | Parent Ptac-PilT | 6.74E-02 | S4 Fig | pilT<sup>K136A</sup> pilU<sup>K137A</sup> pmmB-pilU | 9.52E-04 |
| S1 Fig | Parent Ptac-PilU | 8.85E-02 | S4 Fig | Parent pmmB-pilU | 2.66E-02 |
| S1 Fig | ΔpilT Ptac-PilT | 6.55E-02 | S5 Fig | Parent | 3.12E-01 |
| S1 Fig | ΔpilT Ptac-PilU | 3.13E-06 | S5 Fig | ΔMSHA ΔTCP | 3.73E-01 |
| S1 Fig | pilT<sup>K136A/E204A</sup> ΔpilU Ptac-PilT | 6.32E-02 | S5 Fig | ΔpilTU | 1.59E-05 |
| S1 Fig | pilT<sup>K136A/E204A</sup> ΔpilU Ptac-PilU | 4.05E-02 | S5 Fig | ΔpilTU ΔMSHA ΔTCP | 3.34E-05 |
| S1 Fig | pilT<sup>E204A</sup> ΔpilU Ptac-PilT | 6.02E-02 |  |  |  |
| S1 Fig | pilT<sup>E204A</sup> ΔpilU Ptac-PilU | 7.81E-02 |  |  |  |

**S4 Table.**
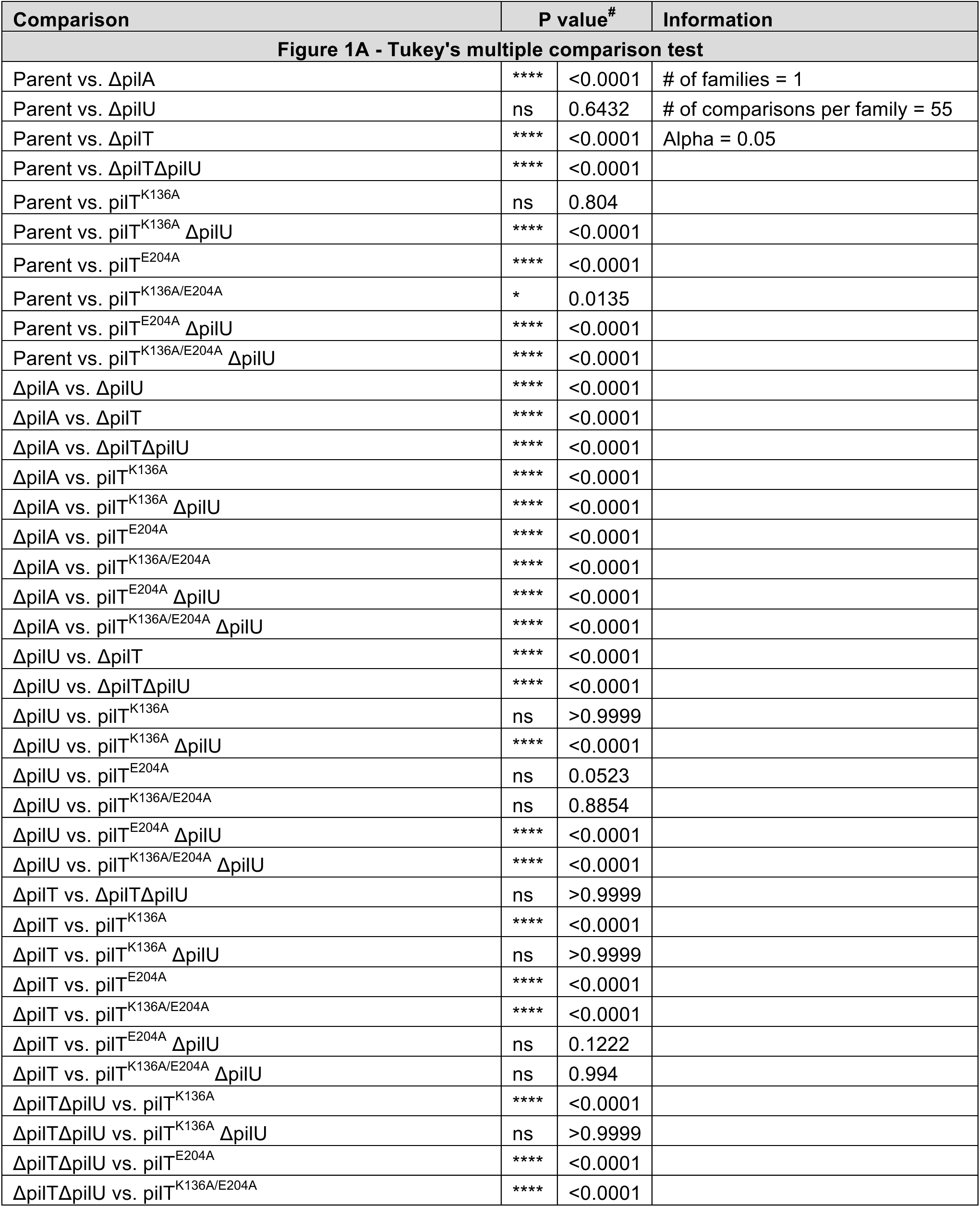
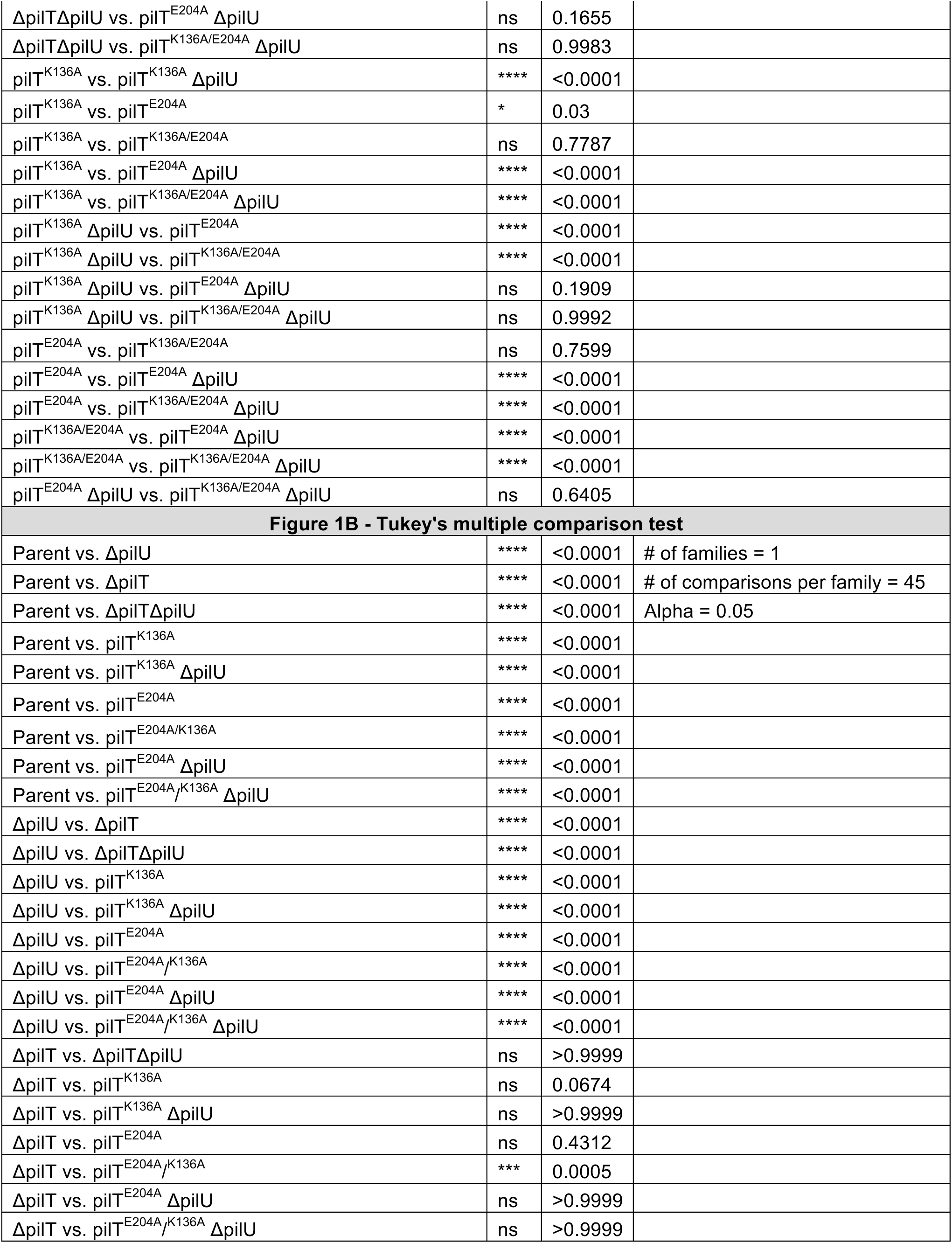
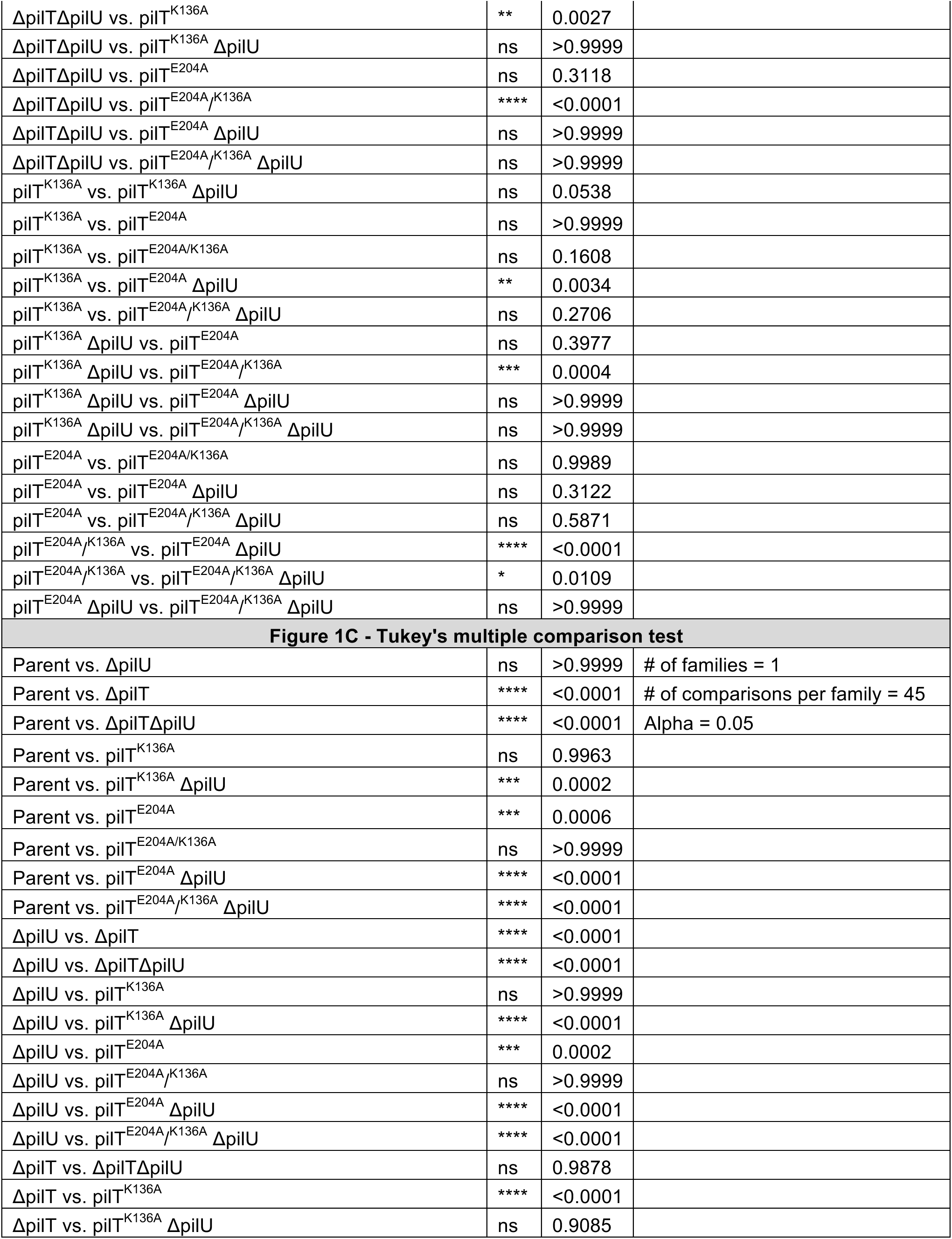
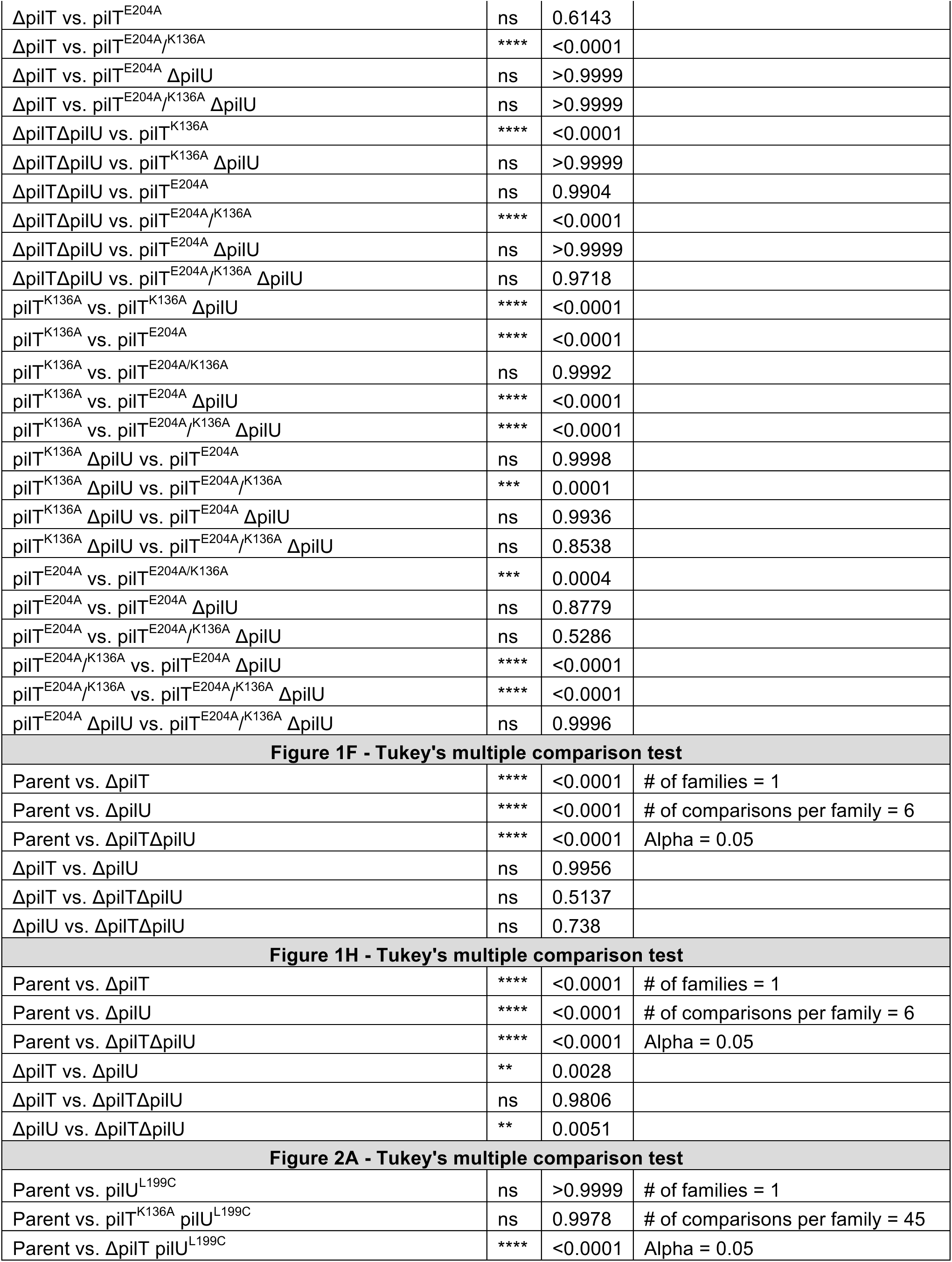
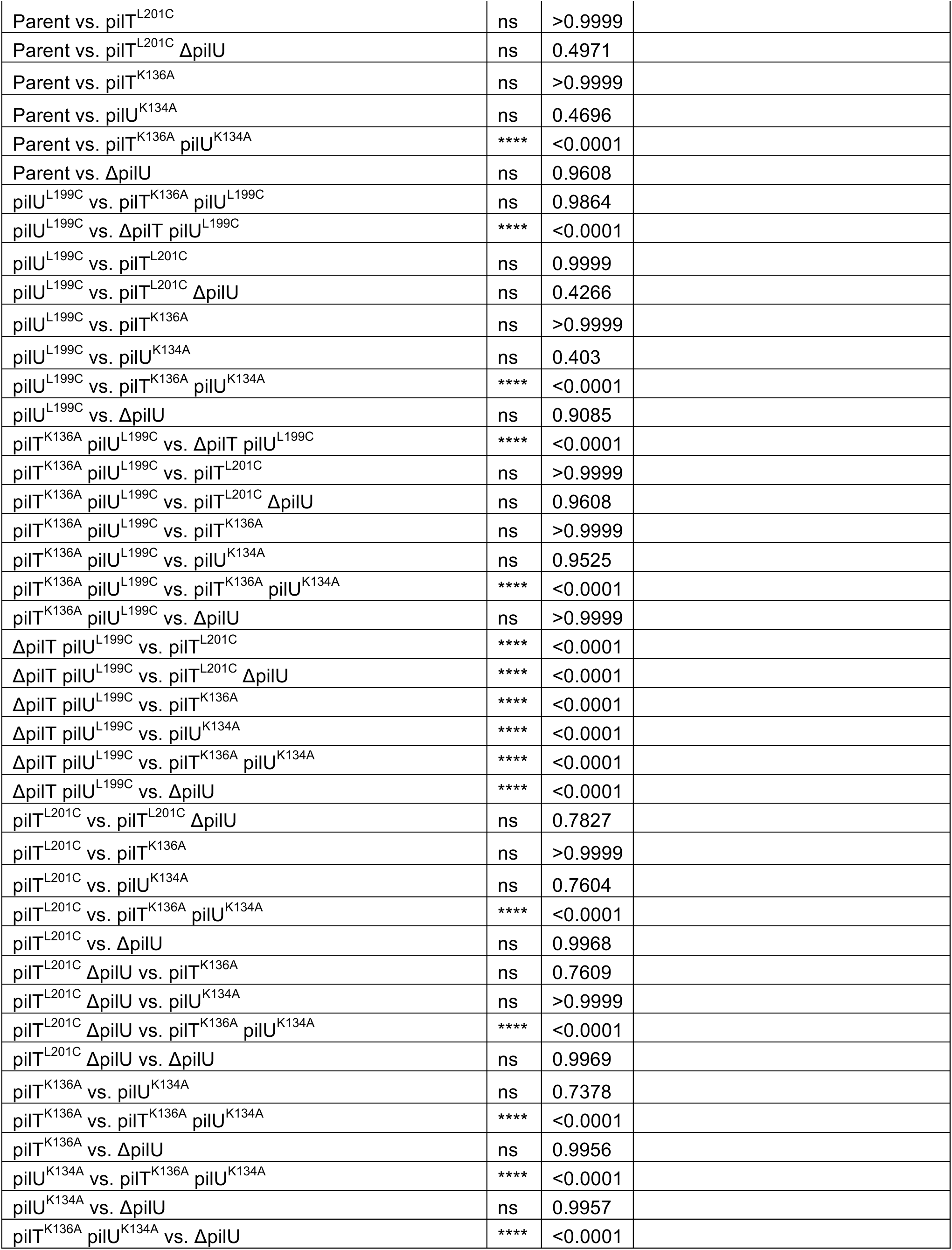
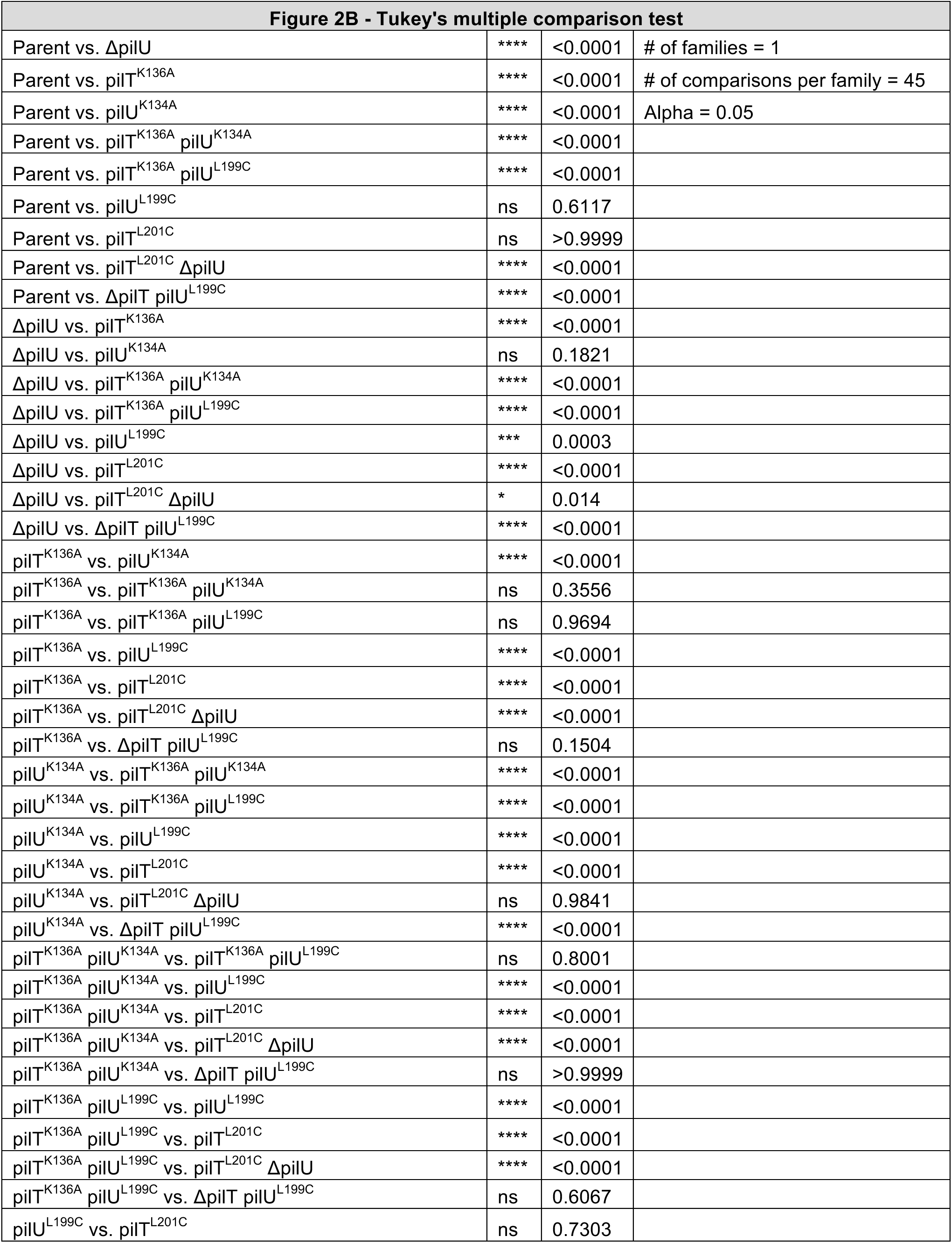
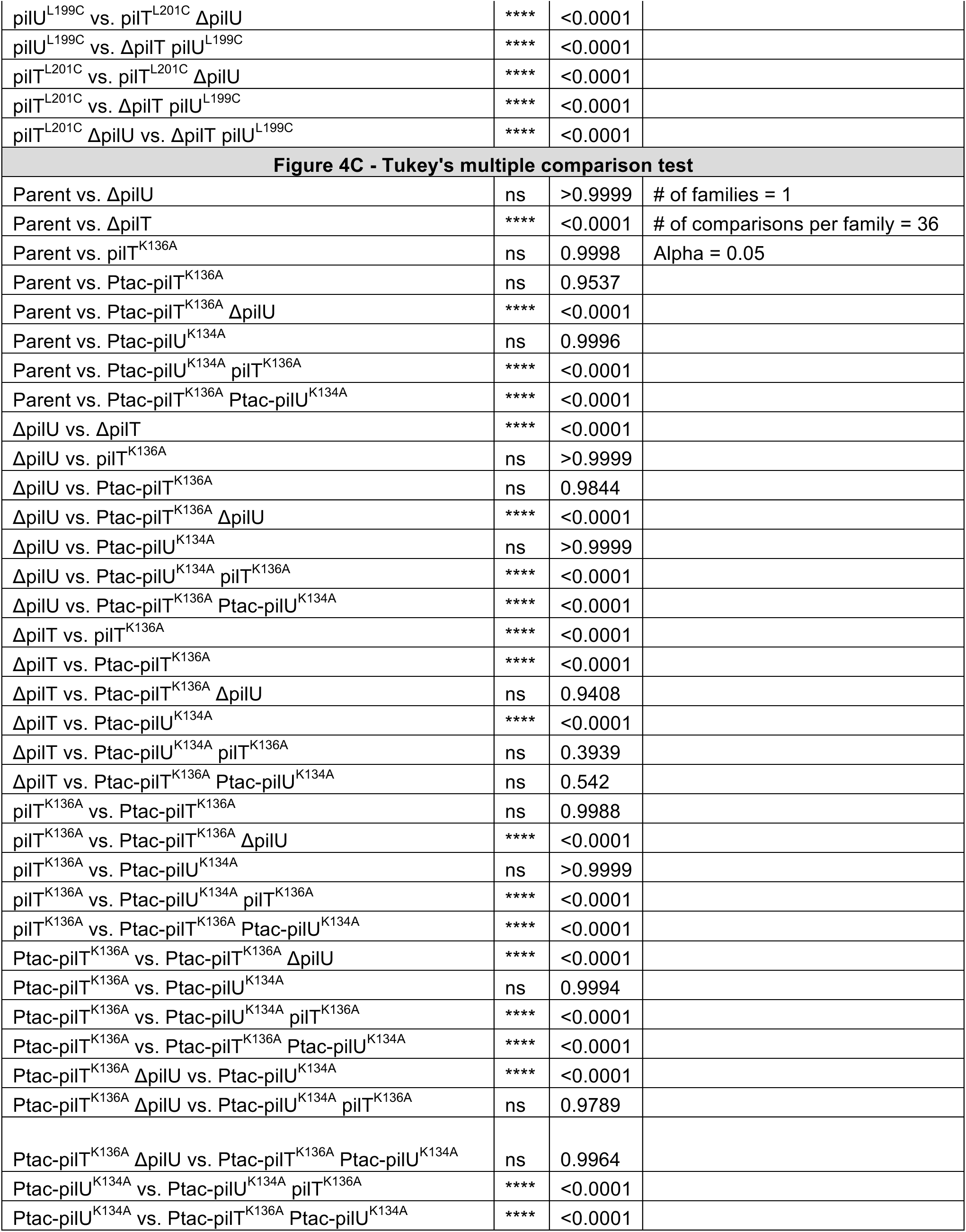
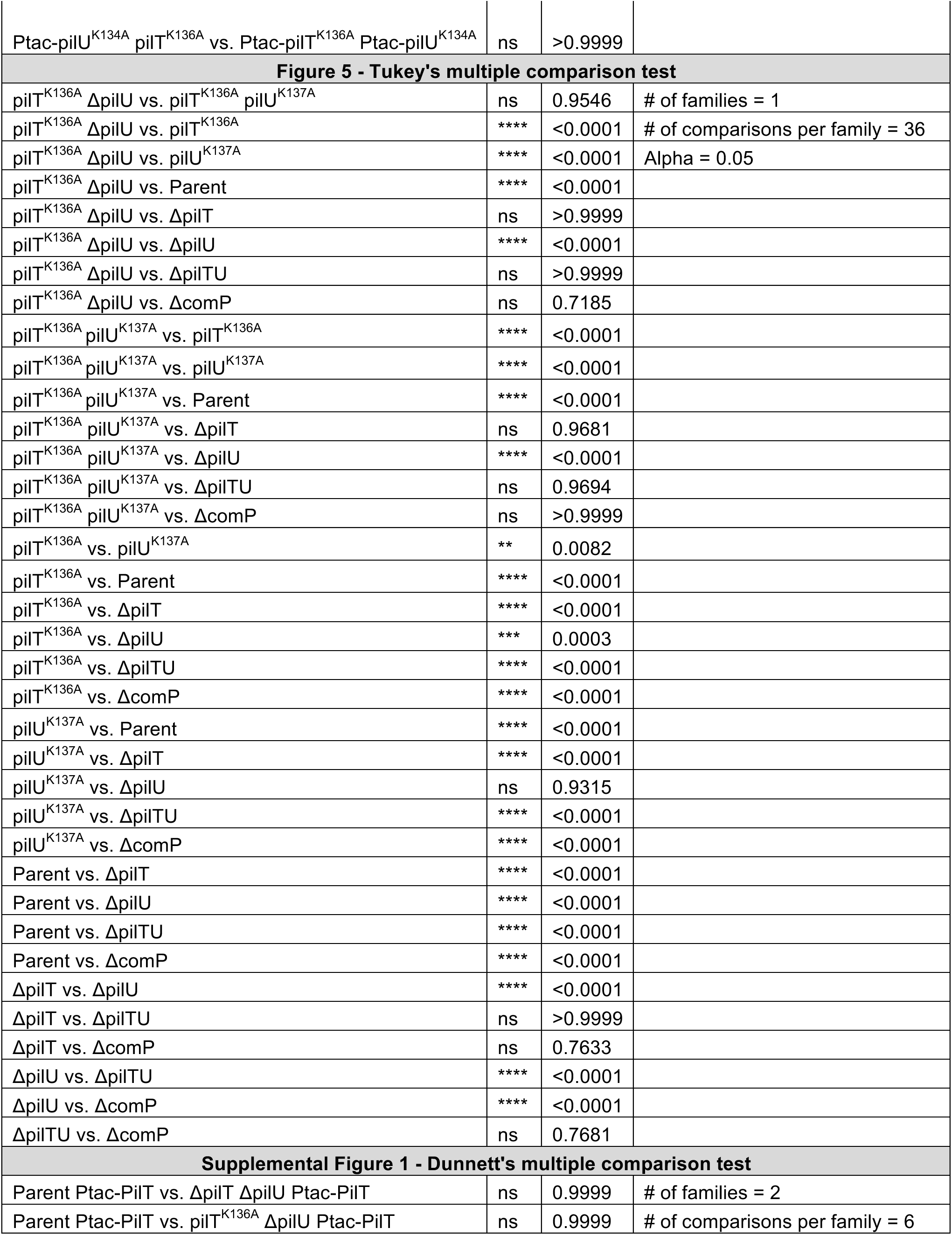
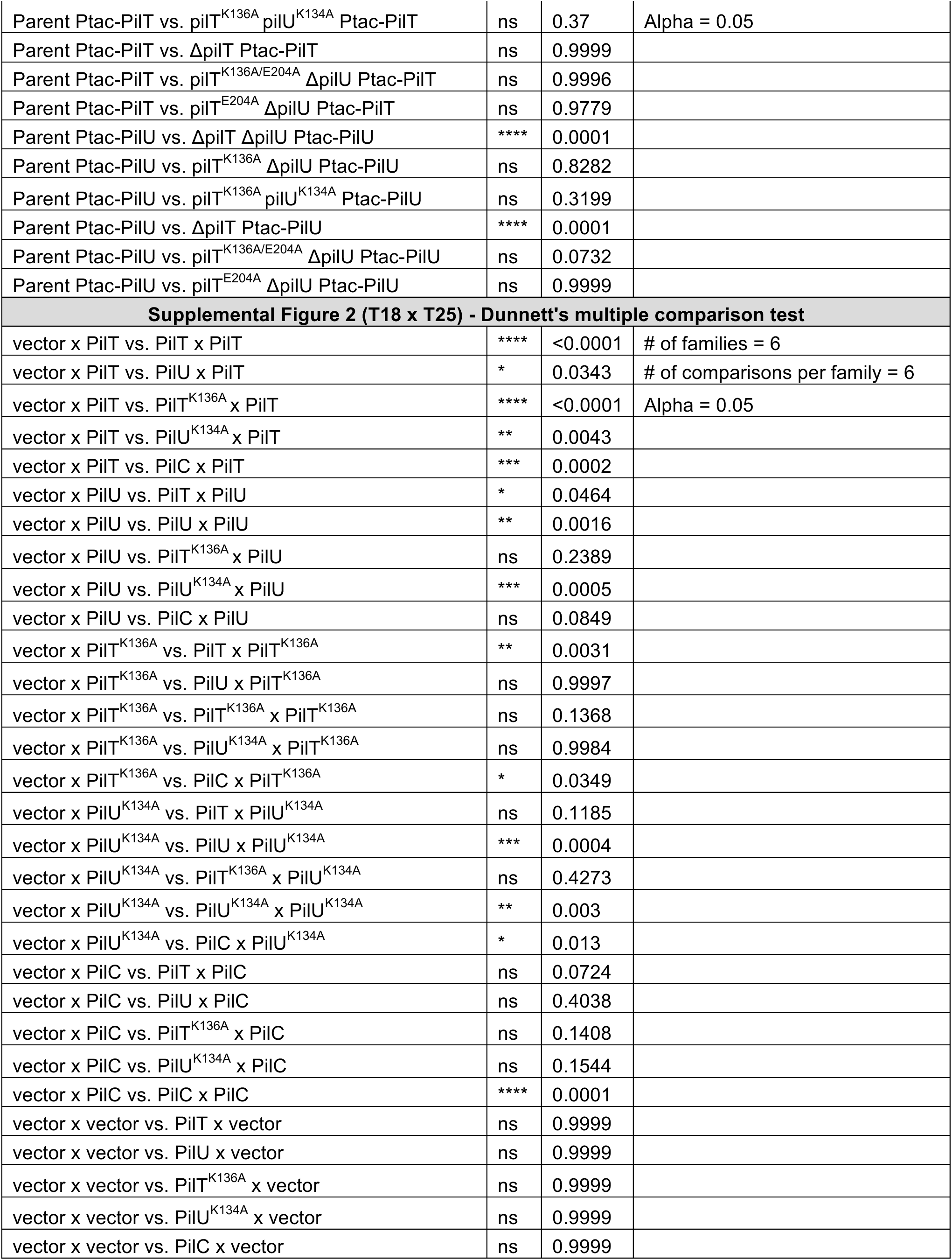
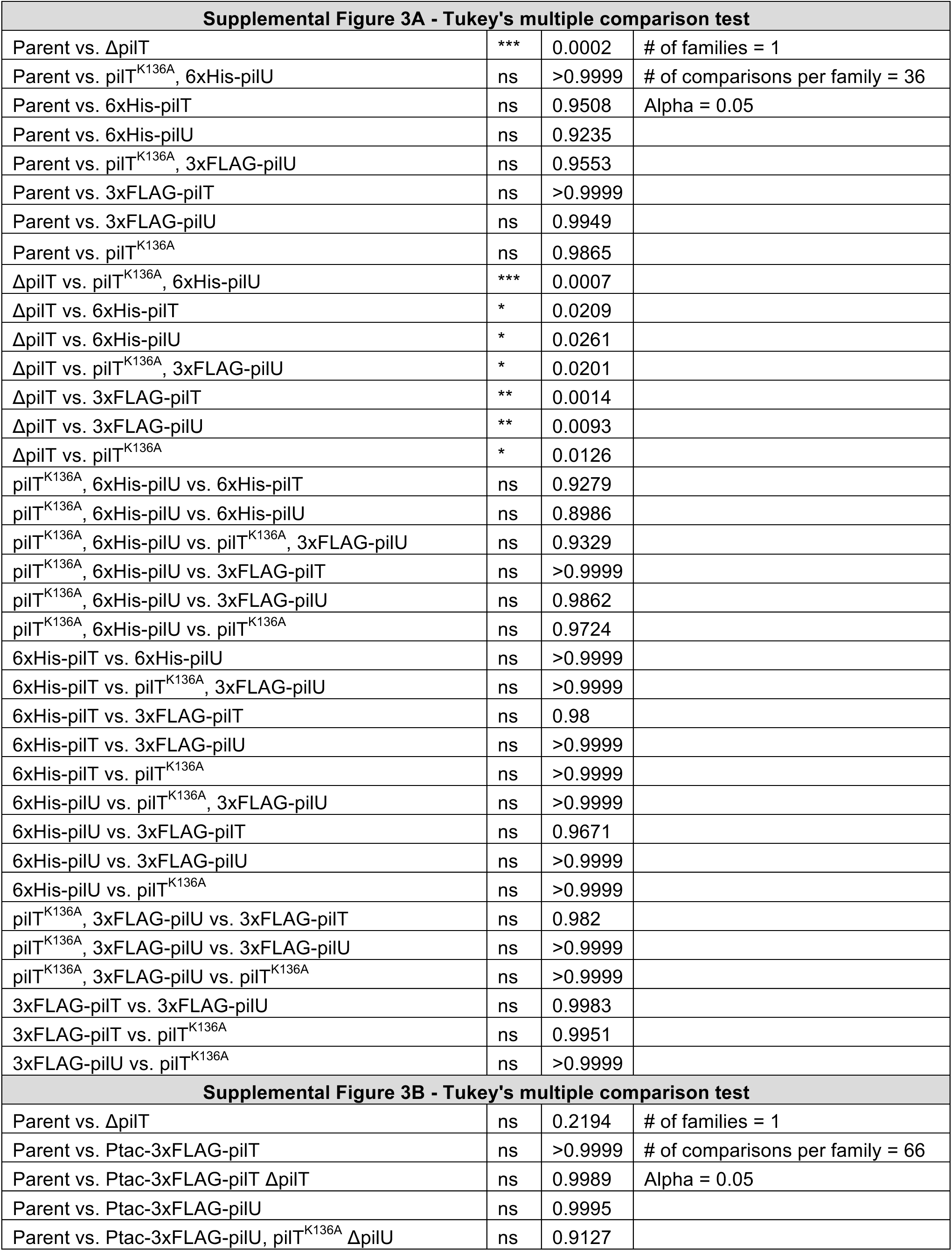
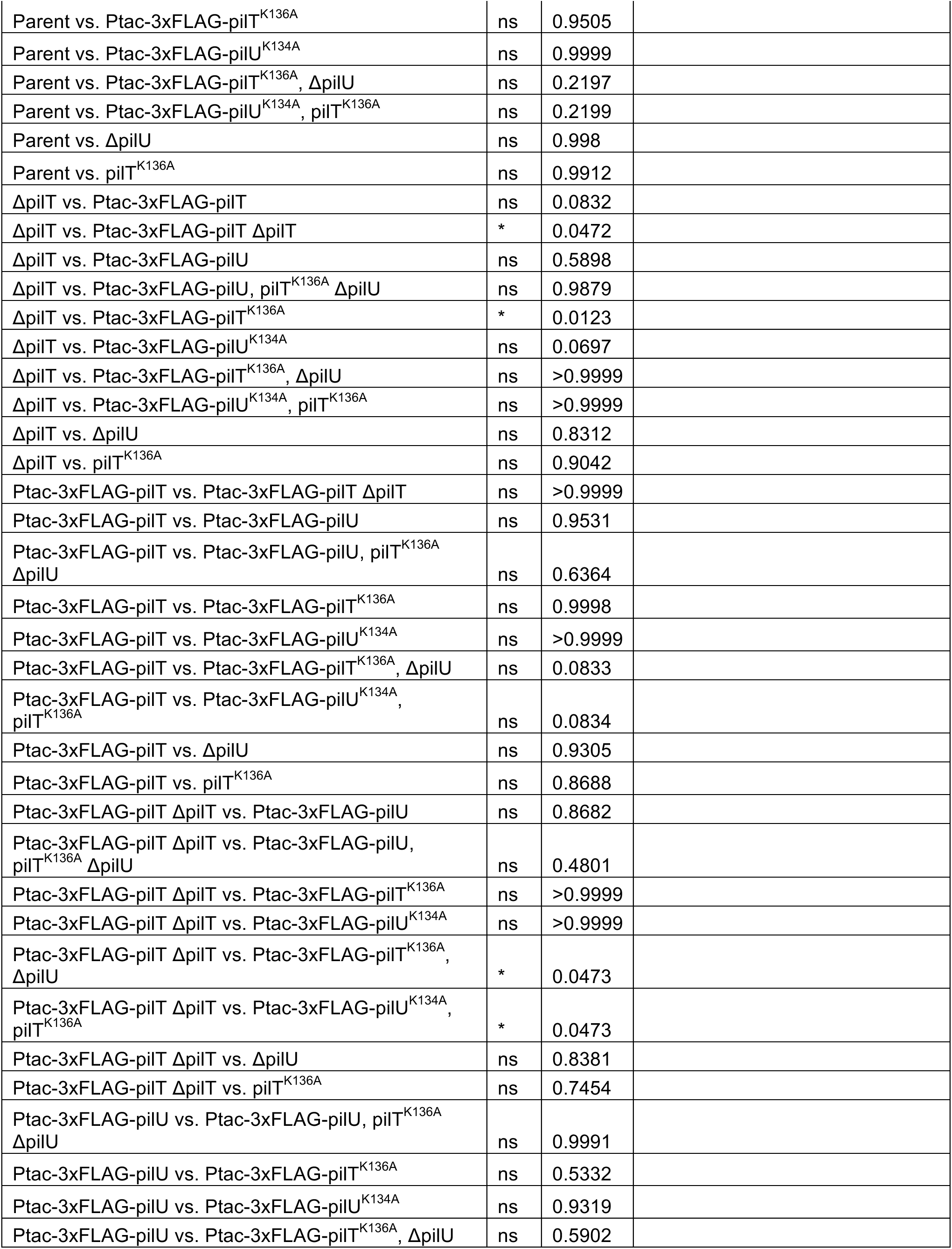
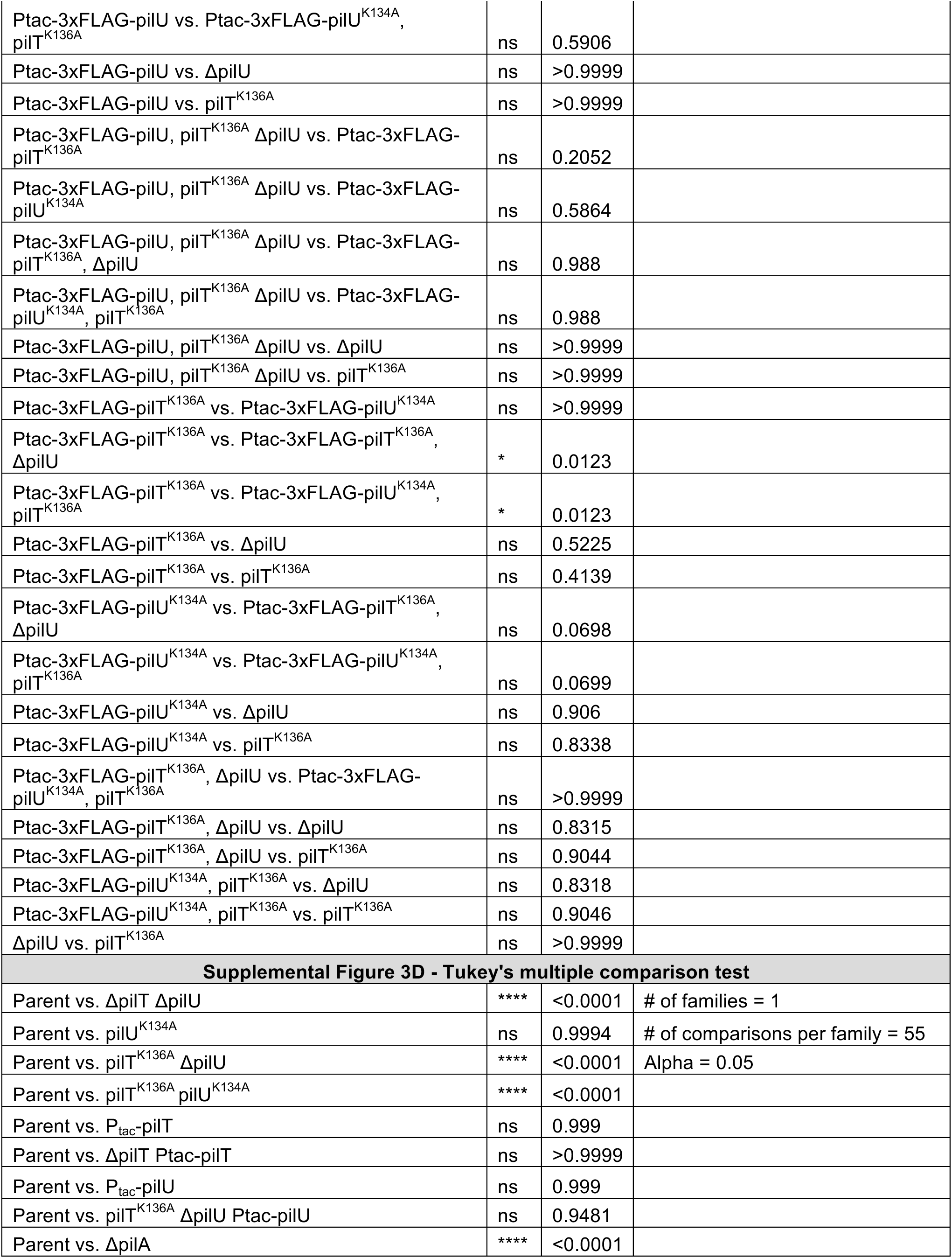
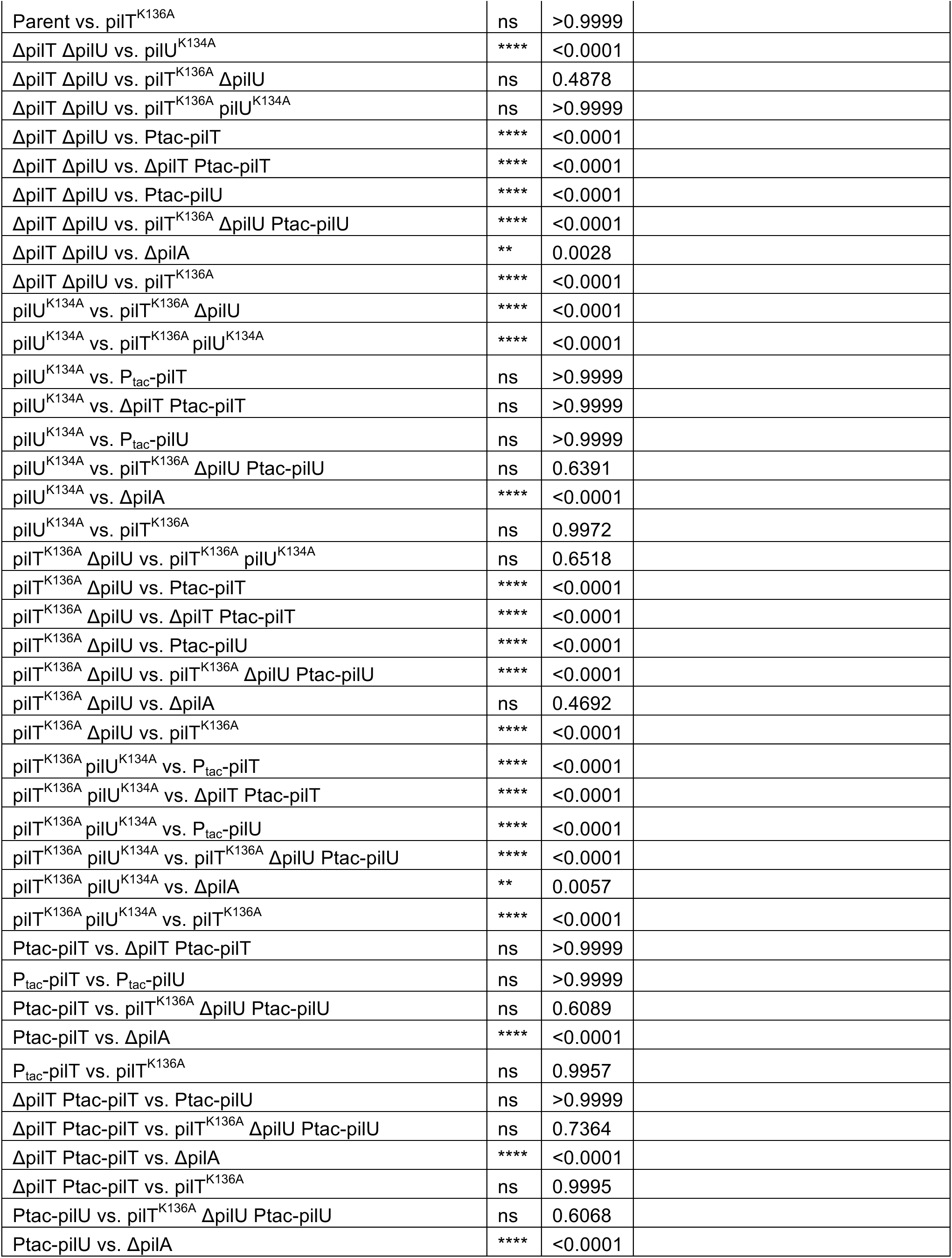
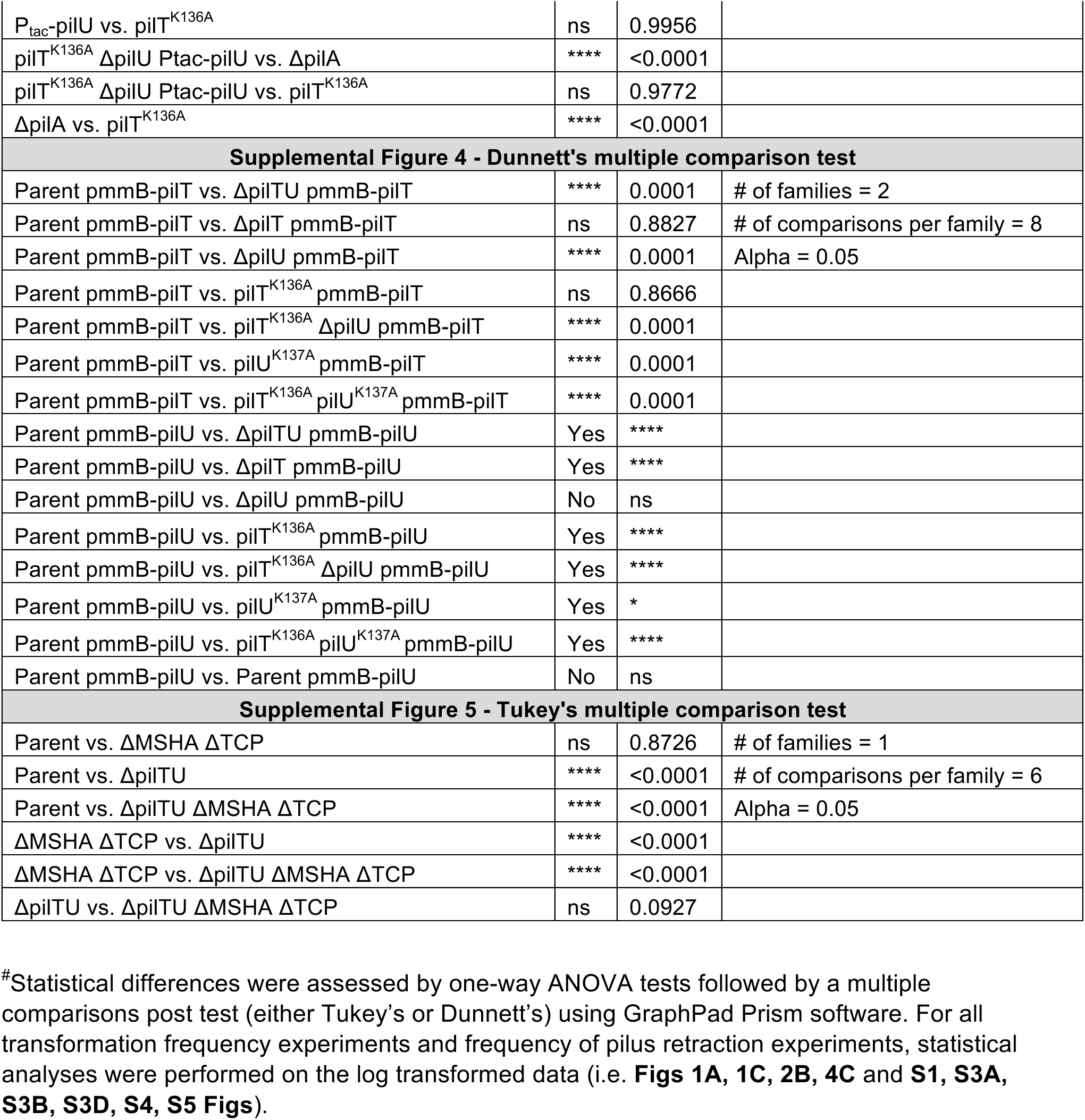
Statistical Comparisons.

